# Coordination through inhibition: control of stabilizing and updating circuits in spatial orientation working memory

**DOI:** 10.1101/819185

**Authors:** Rui Han, Hsuan-Pei Huang, Chia-Lung Chuang, Hung-Hsiu Yen, Wei-Tse Kao, Hui-Yun Chang, Chung-Chuan Lo

## Abstract

Spatial orientation memory plays a crucial role in animal navigation. Recent studies of tethered *Drosophila melanogaster* (fruit fly) in a virtual reality setting showed that the head direction is encoded in the form of an activity bump, i.e. localized neural activity, in the torus-shaped ellipsoid body (EB). However, how this system is involved in orientation working memory is not well understood. We investigated this question using free moving flies (*Drosophila melanogaster*) in a spatial orientation memory task by manipulating two EB subsystems, C and P circuits, which are hypothesized for stabilizing and updating the activity bump, respectively. To this end, we suppressed or activated two types of inhibitory ring neurons (EIP and P) which innervate EB, and we discovered that manipulating the two inhibitory neuron types produced distinct behavioral deficits, suggesting specific roles of the inhibitory neurons in coordinating the stabilization and updating functions of the EB circuits. We further elucidate the neural mechanisms underlying such control circuits using a connectome-constrained spiking neural network model.

**Significance statement:** Head-direction (HD) system has been discovered in rodents for decades. But the detailed neural circuit mechanisms underlying the HD system were only described recently by studies of fruit flies on the similar HD system. However, how this fruit fly HD system involves in orientation memory was not well investigated. The present study addresses this question by investigating free moving flies in a spatial orientation working memory task. By combining neural functional experiments and neural circuit modelling, the study shows how disrupting either of the two subcircuits, one stabilizing and the other updating the neural activity, in the HD system leads to different behavioral impairments. The result suggests specific roles of the HD subcircuits in the spatial orientation working memory.

**Visual Abstract:** 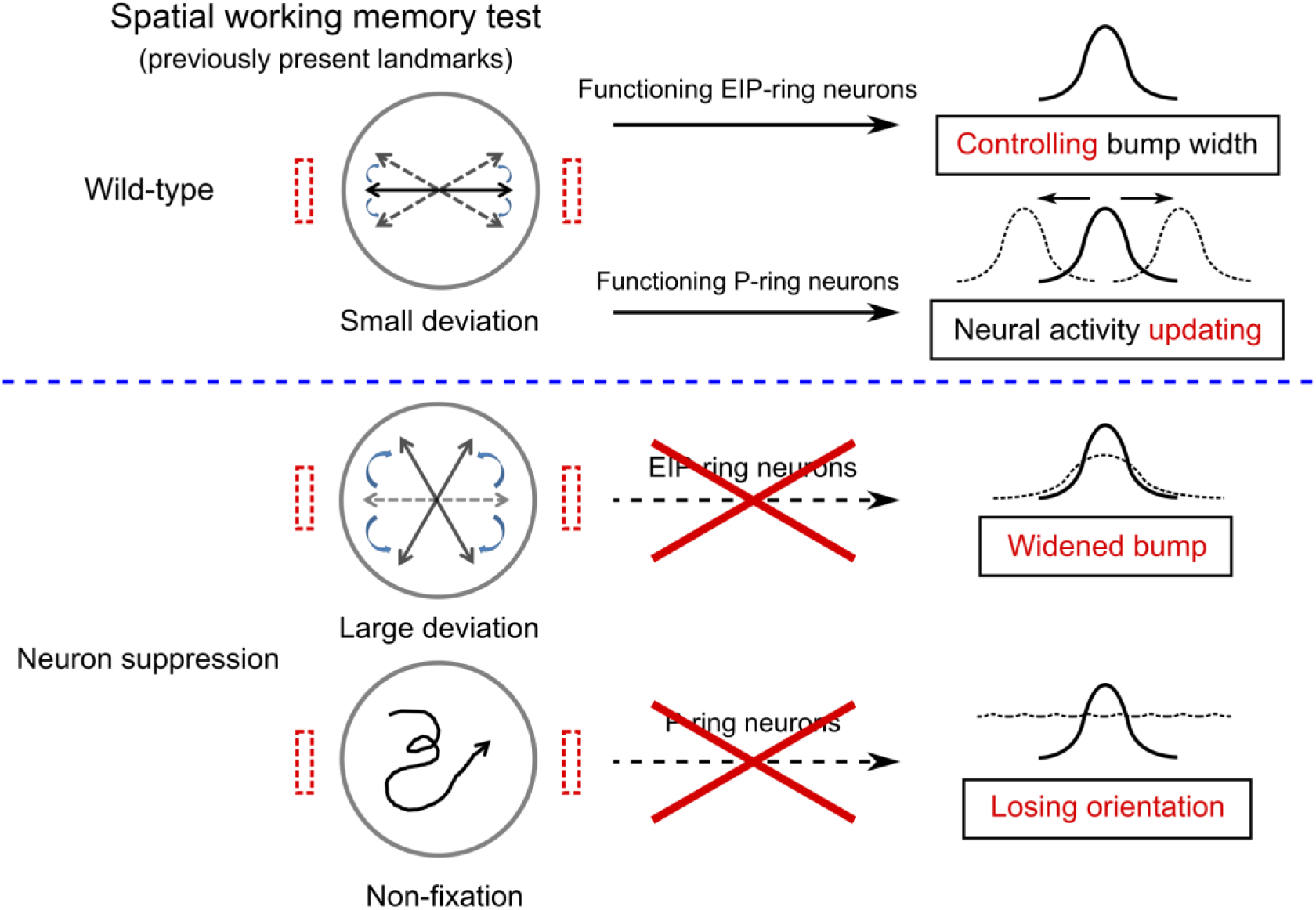

## Introduction

Maintaining spatial orientation is a crucial cognitive capability required for animal navigation (Valerio and Taube, 2016; Yoder and Taube, 2009), and understanding the detailed neural mechanisms of spatial orientation is of great interest to researchers in the fields of neurobiology (Hong et al., 2008; Webb, 2019; Webb and Wystrach, 2016) or neuromorphic engineering (Heisenberg and Wolf, 2013; Lin et al., 2013; Robie et al., 2017). In recent years, significant progress has been made in identifying the neural circuits that support spatial orientation (Dewar et al., 2017) in the central complex of *Drosophila melanogaster* (Strauss, 2002; Turner-Evans and Jayaraman, 2016). The central complex has long been associated with short-term spatial memory, visual pattern memory, and motor control (Liu et al., 2006; Wolf and Heisenberg, 1991). The recent discoveries of head-direction selectivity (Fisher et al., 2019; Kim et al., 2019; Muller et al., 1996; Seelig and Jayaraman, 2015, 2013) and localized neural activity in two central complex neuropils, the ellipsoid body (EB) and the protocerebral bridge (PB), have also linked the central complex to the function of spatial orientation (Dewar et al., 2017; Turner-Evans and Jayaraman, 2016). These studies suggested that the head orientation is encoded by localized neural activity, called activity bump, and the bump location in EB shifts in accordance with changes of heading during movement. The function of the EB neurons resemble that of a compass, and is therefore termed “neural compass” (Clandinin and Giocomo, 2015).

In light of these empirical observations, several neural circuit models of the central complex have been proposed to elucidate the neural circuit mechanisms of head-direction selectivity or other functions associated with the central complex (Cope et al., 2017; Fisher et al., 2019; Givon et al., 2017; Kim et al., 2019, 2017; Pisokas et al., 2020; Stone et al., 2017; Su et al., 2017; Turner-Evans et al., 2017). Some models focused on the stability of the activity bump or on the differences in the circuit dynamics between locus and fruit fly (Kakaria and de Bivort, 2017; Pisokas et al., 2020). Other models studied the plasticity involved in the flexible retinotopic mapping but used simpler firing rate models or schematic models (Fisher et al., 2019; Kim et al., 2019). A large-scale firing-rate neural network model that covered the entire central complex was able to reproduce the steering and homing behavior of bees, but the ellipsoid body circuits were rather simple with minimal details (Stone et al., 2017).

Recently, a spiking-neuron model of the EB-PB circuits was proposed (Su et al., 2017). The model used a more realistic spiking-neuron model and synaptic dynamics to elucidate how the circuits can maintain a stable activity bump when fruit flies switch between forward movement and rotation states in the absence of landmarks. The model suggested the involvement of two subcircuits: one forms an attractor network and maintains (or stabilizes) an activity bump; the other forms a shifter network and shifts (or updates) the bump position in accordance with changes in body orientation. The model successfully demonstrated the angular errors when a fly moved in darkness (Seelig and Jayaraman, 2015) and predicted the asymmetric activity in the PB during rotation (Green et al., 2017).

The model made an important and unique prediction: the function of spatial orientation working memory requires coordinated activation of the bump-maintaining (or stabilizing) and bump-shifting (or updating) circuits that are controlled by the upstream ring neurons.

However, most of the experimental studies used tethered flies in a virtual reality setting and focused on how manipulation of neurons affects the bump activity. It is not clear how these neurons, in particular those involved in stabilizing and updating the activity bump, play roles in cognition-relevant behavior such as spatial orientation memory in free-moving flies with a more realistic behavioral setting. In the present study, we aimed to address these questions and designed a behavioral task of spatial orientation working memory based on the classic Buridan’s paradigm (Götz, 1980; Han et al., 2021; Neuser et al., 2008; Strauss and Pichler, 1998; Yen et al., 2019). Specifically, we manipulated two types of GABAergic ring neurons (Martín-Peña et al., 2014) that are hypothesized to control these neurons. Ring neurons project their axons into EB and inhibit neurons including those display the activity bump. Previous studies have reported the roles of the ring neurons in visually-guided behavior (Ofstad et al., 2011; Pan et al., 2009; Thran et al., 2013), ethanol sensitivity (Awofala, 2011; Kang et al., 2020; Urizar et al., 2007), sleep regulation (Donlea et al., 2018; Guo et al., 2018), olfactory memory (Krashes and Waddell, 2008; Zhang et al., 2013) and mating behavior (Becnel et al., 2011; Ishimoto and Kamikouchi, 2020). However, their roles in the working memory of spatial orientation in the presence or absence of visual cues remain unclear. In addition to the neural functional experiments, we also performed computer simulations using the EB-PB model, which produced neural activities that were consistent with the behavioral changes observed in the fruit flies with different experimental conditions. The present study provides a detailed picture on how coordinated activation between the neural processes of stabilization and update plays a crucial role in spatial orientation working memory.

## Materials and Methods

### Fly strains

In the present study we used both male and female flies. Flies were raised at 25*°*C with a 12:12 h light-dark cycle with a humidity level of ∼50%. Wild-type and transgenic flies were obtained from the Bloomington *Drosophila* Stock Center, *Drosophila* Genomics Resources Center and Brain Research Center (BRC) in National Tsing Hua University (NTHU). We used wild-type *w^+^* (Brain Research Center, National Tsing Hua University), *;; UAS-Kir2.1* (Guo et al., 2015; Shuai et al., 2015) (Brain Research Center, National Tsing Hua University), *;;VT5404-GAL4* (Lin et al., 2013) (Brain Research Center, National Tsing Hua University), *;;UAS-CsChrimson.mVenus* (Bloomington Stock number #55136), *;;UAS-GFP* (Bloomington Stock number #1522), *tub-GAL80^ts^;;* (Bloomington Stock number #7017), *c105-GAL4;;* (Humberg et al., 2018) (Bloomington Stock number #30822), *;;UAS-TNT* (Eisel et al., 1993; Sweeney et al., 1995) (Bloomington Stock number #28997), *;;ninaE* (*Drosophila* Genomics Resources Center number #109599). For the neuron suppression experiments, all GAL4 lines were crossed to the *UAS-Kir2.1* or *UAS-TNT* effector lines and the expression can be controlled by temperature with the combination of *tub-GAL80^ts^*. For the optogenetic experiments, all *GAL4* lines were crossed to the *UAS-CsChrimson.mVenus* effector lines. Further information and requests for resources and reagents should be directed to and will be fulfilled by the corresponding author.

### The arena

We conducted the behavioral tasks with flies in a circular arena constructed in house. We followed the general design principles described in a previous study (Yen et al., 2019). The arena used in the present study had a central circular platform of 85 mm in diameter and was surrounded by a 360*°* LED display with water filling the basin surrounding the platform (Fig. 1A). The display was made of 20 LED panels and each panel consists of four 8 × 8 LED matrices (Small 1.2’’ 8 × 8 Ultra Bright Yellow-Green LED Matrix KWM-30881CUGB). The entire LED screen measured 200 mm in diameter and 130 mm in height. We used green LEDs with a wavelength of 572 nm, which is close to the peak sensitivity of fly eyes (Longden, 2016). Each vertical line of the LEDs could be individually controlled by a personal computer via an Arduino board (Arduino Shield MEGA2560). A CCD camera was mounted directly above the center of the platform and was used to record the movement trajectories of the flies using a Python script developed in house.

**Figure 1.**
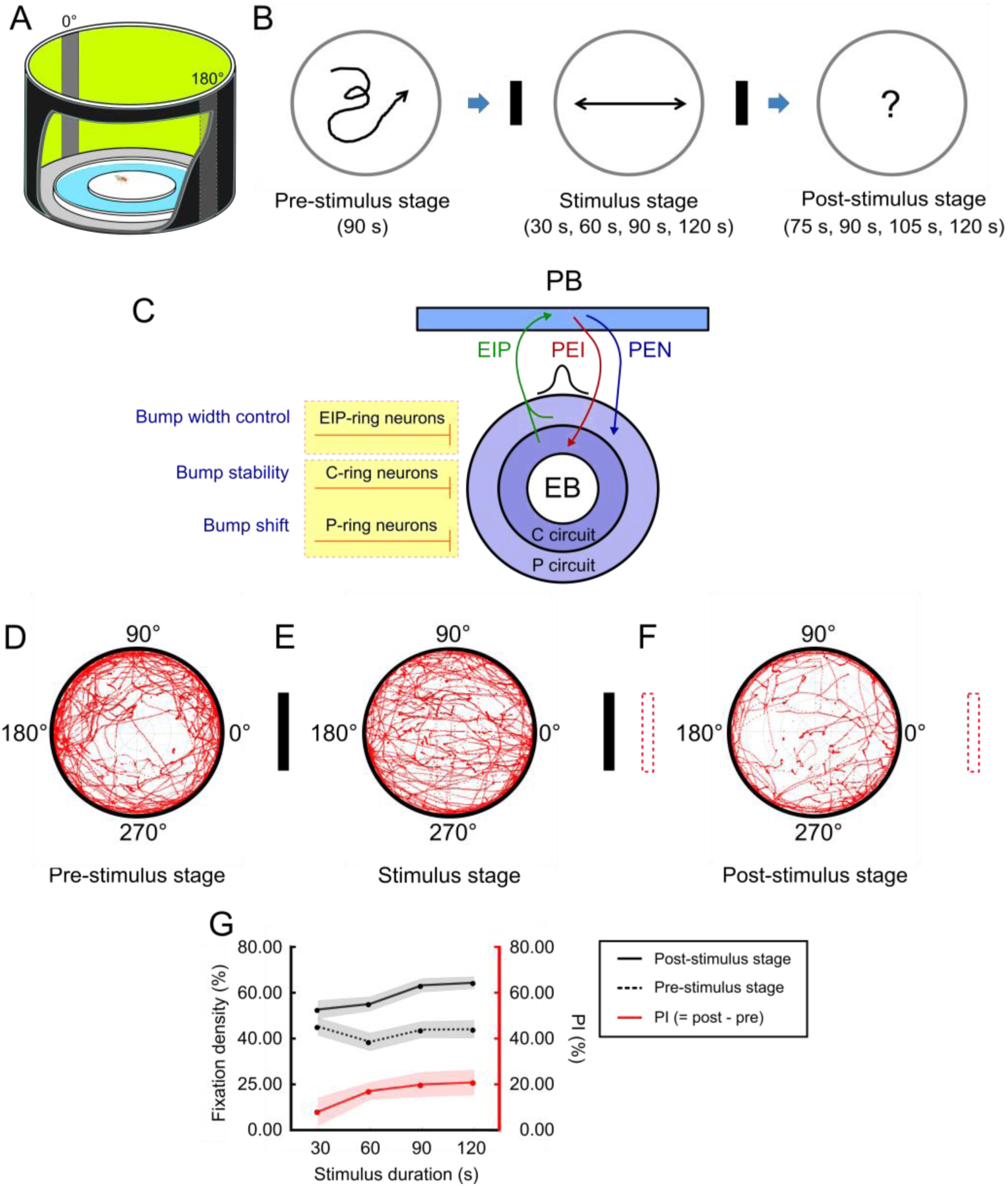
The model and experiment of spatial orientation working memory for *Drosophila melanogaster* (fruit fly). The performance of the wild-type flies indicated strong working memory of the landmark directions. ***A***, The behavior arena featured a central platform surrounded by water and by a 360*°* LED screen that displayed two vertical black strips as the visual landmarks at 0*°* and 180*°*. ***B***, The task, designed based on Buridan’s paradigm (Götz, 1980; Neuser et al., 2008; Strauss and Pichler, 1998), was divided into three stages: The pre-stimulus stage (90 s), stimulus stage (variable duration), and post-stimulus stage (variable duration). In each trial, a single fruit fly was placed at the center of the platform and was allowed to freely move on the platform in the whole trial. ***C*,** Schematic diagram of the EB-PB neural network (Su et al., 2017). The EIP, PEI and PEN neurons support the activity bump which encodes the head direction with respect to a cued location. The bump is modulated by three types of GABAergic ring neurons, EIP-ring, P-ring and C-ring neurons, and each performs different functions. These ring neurons can be targeted by specific *GAL4* drivers as shown in Extended Data Figure 1-1. ***D-F***, The walking patterns as indicated by the red trajectories of the flies (n = 30) in the pre-stimulus, stimulus, and post-stimulus stages. In the stimulus stage, there was a clear pattern of fixation toward the landmarks. The fixation behavior was reduced, but still statistically significant in the post-stimulus stage. The definition of the movement direction and the trajectory distribution are shown in Extended Data Figure 1-2. ***G***, The fixation density (fixation duration / stage duration) of the post-stimulus stage (black solid), the pre-stimulus stage (black dashed) and their differences, termed performance index (*PI*) (red) as functions of the stimulus exposure duration. The strength of memory, which is indicated by *PI*, increases with the stimulus exposure duration. Schematics of neural mechanisms underlying spatial orientation memory and its circuit model are show in Extended Data Figure 1-3. Movement patterns of individual flies are displayed in Extended Data Figure 1-4.

### The *GAL4* drivers

We used *c105-GAL4* line to target the EIP-ring neurons and *VT5404-GAL4* to target the P-ring neurons in the present study (Extended Data Fig. 1-1A). We have also inspected other EIP-ring neuron expressed drivers including *R31A12-GAL4* (Omoto et al., 2018), *VT039763-GAL4*, and *VT39763-GAL4* (Lin et al., 2013) (Extended Data Fig. 1-1B). For P-ring neuron expressed drivers, we have inspected *VT005404-GAL4* and *R14G09-GAL4* (Omoto et al., 2018) (Extended Data Fig. 1-1C). C-ring neuron expressed driver *VT011965-GAL4* (Omoto et al., 2018) are inspected as well (Extended Data Fig. 1-1D). However, these drivers are less specific and are expressed in many other brain regions. Specificity of the drivers is crucial to the present study as it involves behavioral experiments. Therefore, we only used *c105-GAL4* and *VT5404-GAL4* in the present study.

### The spatial orientation memory task

We used 3-5 day-old flies and clipped the wings 1 day before the experiments (Neuser et al., 2008). In addition to wild-type flies (genotype: *w^+^*), we used flies with suppressed EIP-ring or P-ring neurons, which was achieved by hyperpolarizing the neurons (32*°*C, *c105-GAL4, tub-GAL80^ts^;; UAS-Kir2.1* for EIP-ring neuron suppression and *;;VT5404-GAL4, tub-GAL80^ts^ / UAS-Kir2.1* for P-ring neuron suppression) or by blocking the neurotransmitters (32 *°* C, *c105-GAL4, tub-GAL80^ts^;; UAS-TNT* for EIP-ring neuron suppression and *;;VT5404-GAL4, tub-GAL80^ts^ / UAS-TNT* for P-ring neuron suppression).

After the wings were clipped, the flies were placed in an 18*°*C incubator in order for *GAL80^ts^* to bind *GAL4* and inhibited the transcription activity of the *GAL4* (control groups), or in a 32*°*C incubator to relieve the inhibition of *GAL4* proteins so that they can drive the expression of *UAS* (experimental groups) (KaiXia et al., 2016) for 1 day. All experiments were performed between 10am and 5pm. We also used optogenetics to transiently activate EIP-ring or P-ring neurons. The expression of the effector (*UAS-CsChrimson.mVenus*) was controlled by red lights (625 nm) exposure after feeding all-*trans*-retinal (100 *µm*) in a dark environment for 7 days (Wu et al., 2014).

Based on the protocol proposed in an earlier study (Yen et al., 2019), the spatial orientation memory task consisted of three stages: The pre-stimulus stage, stimulus stage, and post-stimulus stage (Fig. 1B). Only one fly was submitted to the task in each trial and the fly was allowed to freely move on the circular platform. The pre-stimulus stage lasted 90 s, during which all LEDs were turned on and no visual landmark was presented on the screen. During the stimulus stage, two vertical black strips, each 30*°* wide and separated by 180*°*, were presented on the screen as visual landmarks. We tested four different durations (30, 60, 90 and 120 s) for the stimulus stage with wild-type flies, and used 60 s in the subsequent neural functional experiments. In the post-stimulus stage, the two landmarks were removed and all LEDs were turned on; this stimulus condition was identical to that used in the pre-stimulus stage. The duration of the post-stimulus stage was 75, 90, 105, and 120 s for the four different durations, 30, 60, 90, and 120 s, of the stimulus stage, respectively. The movement trajectories of the flies were recorded using a CCD camera at a speed of 20-25 frames per second. The position of the fly in each frame was captured by a Python script and saved for post-experiment analysis.

### Optogenetic activation

Red LED array made up of 96 LEDs (625 nm, RED LED SMD 5050) were used to activate *CsChrimson* during the behavioral task. The LED array was placed below the platform in the arena. The platform was made of acrylonitrile-butadiene-styrene (ABS) and was partially transparent to light due to its 1mm of thickness. We used two protocols of photo activation. In the first protocol, the red light was turned on during the last 30 s of the stimulus stage. In the second protocol, the red light was turned on for only 10 s starting at 20 s after the post-stimulus stage began.

### Confocal images

7-day-old adult flies brains were fixed, mounted in phosphate buffer saline (PBS) as previously described (Chang and Ready, 2000). Images were scanned with Zeiss LSM 510 confocal microscopy.

### Data analysis

One of the goals of the present study was to investigate the alternation of spatial orientation memory, as indicated by the movement direction, under various neural manipulations. Therefore, we designed behavioral measures and analytical methods for this purpose. We first defined the performance index (*PI*) that quantifies how well the flies fixed on the cued locations against a reference direction, i.e. the perpendicular direction. This single-valued measure is easy to be analyzed and compared statistically. However, we discovered that in some conditions, the flies exhibited strong fixation behavior but on the wrong directions. This behavior has a different implication from that exhibiting no fixation. *PI* cannot distinguish the two different behaviors and therefore we designed the radar plot to capture the differences. We present the two analyses for all spatial orientation tasks conducted in the present study and the detailed definitions of the two measures are described below.

To analyze the movement direction of each fly (Liske, 1977) in each video frame, we first calculated the speed vector based on the difference in the coordinates of the fly between two consecutive frames. The movement direction was represented by the degree value, *θ*, on the screen at the point that the speed vector projected to (Extended Data Fig.1-2). For better presentation and analysis, we calculated the percentages, *p*(*θ*), of the movement direction in each of the 12 quantiles in 360 ° . *θ* represented the degree corresponding to the center of the quantile. Each quantile spanned 30° on the screen. *p*(*θ*) was calculated every 5 s with a 15 s sliding time window. Fixation behavior was characterized by a significantly higher percentage of the movement direction falling within ±15° of 0° and 180° on the screen, the quantiles that the two visual landmarks occupied.

To further quantify the performance of the flies in terms of the spatial orientation memory, we defined the performance index (*PI*). We first calculated *P*(0°; 180°) = *p*(0°) + *p* (180°) in each stage for each fly. Next, for a given stage, we calculated the fixation density *FD*(0°; 180°), which was defined as the number of 5 s epochs in which *P*(0°; 180°) > 1/6 (or 16.67%) in a stage divided by the total number of epochs of that stage. 16.67% was the expected percentage that a fly spent in two quantiles, e.g. 0° and 180°, if it moved randomly. Due to the highly variable nature of the fly movement, *P* could be larger than 16.67% in multiple different directions in an epoch. This led to a non-zero *FD* in more than one direction, even if the fly performed the random walk. Therefore, we stress that measuring *FD* along in the third stage did not provide correct information about the presence of memory, which should be evaluated by calculating the difference in *FD*(0°; 180°) between the third and the first stage. This is given by *PI*, which was defined as

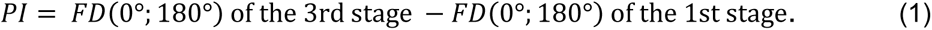

In addition to *PI*, which only measured the fixation behavior at the 0° and 180°, we also visualized the fixation behavior at all directions by the radar plots, which indicated the frequencies *p*(*θ*) of the movement direction in each quantile. In the plot, each pair of quantiles that were 180° apart, e.g. quantiles centered at 90*°* and -90*°*, was represented by the same value, which was their averaged percentage. This made the radar plot point-asymmetric. If a fly did not exhibit any fixation behavior, it would move toward any direction with equal probability due to the point symmetric property of the arena, and the expected value of each pair of quantiles on the plot was 1/6 (∼16.67%). A value that was significantly different from 1/6 indicates some form of fixation behavior.

The radar plots allowed us to visually inspect the existence of fixation behavior and the main direction that the flies fixated on with respect to the directions of the visual landmarks. To further quantify these two behavioral properties, we calculated the fixation deviation angle (*σ_f_*) and the fixation strength ( *F* ). The fixation deviation angle ( *σ_f_* ) represented the direction of the strongest fixation tendency and was defined by the angle *σ*, which yielded the smallest second moments *M*(*σ*) on the radar plot:

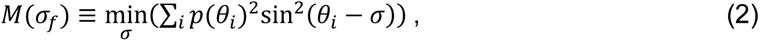

where *θ_i_* is the angle of each quantile (0*°*, 30*°*, 60*°*, 90*°*, ….). *σ_f_* can be obtained by taking the derivative of *M*(*σ*) with respect to *σ*.

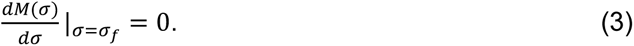

Plugging *θ_i_* and *p*(*θ_i_*) into this equation and solving for *σ* using trigonometric identities, we obtain

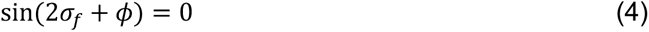

and

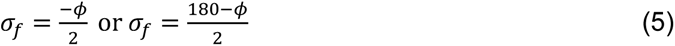

where *Φ* is a variable depending on *p*(*θ_i_*) based on the following equations:

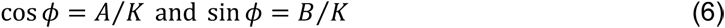

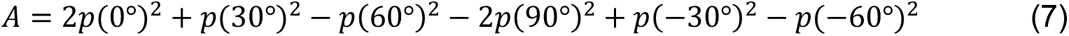

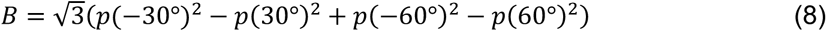

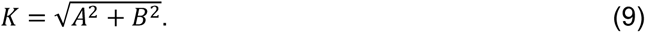

The deviation angle given in Equation (6) corresponds to two values. One gives rise to the minimum value of *M* (denoted as *M_min_*) and the other gives rise to the maximum value of *M* (denoted as *M_max_*). Finally, the fixation strength (*FS*) is defined as follows:

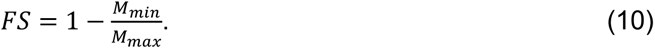

To determine what level of *FS* indicates a significant fixation, we analyzed the *FS* for all flies in the control groups, during the pre-stimulus stage. The average *FS* was 0.0544, and the standard deviation was 0.0273. We took two standard deviations above the mean (*FS* = 0.1090) as our statistical criterion for the fixation behavior.

We used the performance index (*PI*) and the fixation strength (*FS*) to quantify the fixation behavior. Although they seem to be redundant, they serve different purposes. The *PI* specifically measures the difference in the fixation (on the cued directions) between two stages and is an indicator of the spatial orientation memory if *PI* is measured for the third and first stages. *PI* does not measure whether the flies also fixate on the other directions. By contrast, the *FS* measures the fixation toward any direction. The exact azimuth angle of the fixation is specified by *σ_f_*. *FS* is useful if we want to investigate the deviation of fixation from the cued directions.

### Analysis of locomotion

To test whether the genetic manipulation in the central complex led to a deficit in locomotion, which in turn contributed to the change in fixation behavior, we analysed the average movement speed and activity level of the flies. The average speed was calculated by dividing the total movement distance by the total duration of movement in one stage of the task. The activity level was defined as the percentage of video frames in which the fly moved.

### Vision test

We tested the vision of the flies to ensure that the observed behavioral changes were not due to vision impairment. In the first stage of the test, we allowed a fly to freely move on the platform for 60 s. In the second stage, we put a laser spot (50 mW, wavelength = 532 nm) on the platform and gradually moved the spot toward the fly until the laser hit the fly’s body. We repeated this procedure several times. Due to its power, the laser spot was strongly aversive to the flies, and they quickly learned to avoid the laser spot. Typically, after a couple of hits, the flies began to escape the approaching laser spot before it reached them. As a control, we also tested flies with deficient photosensors (genotype: *;;ninaE*) (Movie 1) and wild-type flies with amputation of the foreleg (genotype: *w^+^*) (Movie 2) (Isakov et al., 2016). Such escape behavior would not occur if the flies could not visually perceive the approaching laser spot. We measured the escape rate, *r*, which was defined as:

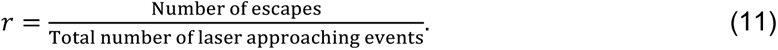

We then compared *r* between the transgenic flies and the wild-type flies. We would expect a lower *r* for the flies if they were visually impaired.

### Statistical analysis

For wild-type flies, 10-25 individuals were used for each experimental condition in the short-term orientation memory test. For the neural functional experiments and optogenetic activation experiments, 9-20 flies were used in each group.

The statistical analyses were performed using Statistical Product and Service Solutions 22.0 (SPSS 22.0). The performance index was analysed using the multi-factor within group analysis of variance, and the differences between groups with different genetic conditions were evaluated using the mixed variance analysis.

### The EB-PB circuit model

We used a previously proposed spiking neural network model of partial central complex (Su et al., 2017), which was built based on the connectomic data of the EB and PB (Lin et al., 2013; Wolff et al., 2015). The model was simulated using the Flysim simulator (Huang et al., 2019). The detailed model equations, neuronal connectivity and model parameters are available in Su et al., 2017. Here we only provide a concise description of the model. The model network consists of two coupled ring circuits with attractor dynamics (Fig. 1C, Extended Data Fig. 1-3A-F): (1) The recurrent C circuit (Extended Data Fig. 1-3G), formed by EIP (or E-PG) and PEI, is responsible for maintaining a stationary activity bump when a fruit fly moves straight, or when it stops. (2) The shifter P circuit (Extended Data Fig. 1-3H), formed by EIP and PEN (or P-EN) neurons (Green et al., 2019; Kim et al., 2017; Turner-Evans et al., 2017), is responsible for shifting/updating the bump in accordance with the body rotation or the motion of a salient visual object so that the bump always indicates the correct head direction with respective to a cued location. Each of the ring circuits is mapped topographically to 360° of the horizontal plan of the external space. The P circuit can be divided into two sub-circuits and each corresponds to a unilateral PB. Activation of each sub-circuit, or one side of PB, shifts the bump in different directions. The model predicts that alternated activation of the C and P circuits is crucial for a fruit fly because its movement pattern is usually characterized by interleaved straight movement and rotation. The model further predicts that the alternated activation of the C and P circuits is controlled by corresponding GABAergic C-ring and P-ring neurons, which inhibit PEI and PEN neurons, respectively (Extended Data Fig. 1-3I). The C-ring neurons regulate the stability of the bump while the P-ring neurons regulate the bump shift. There is an additional type of ring neurons (EIP-ring neurons), which always activate in order to provide global inhibition that maintains a narrow bump (Extended Data Fig. 1-3H). Therefore, to validate the hypotheses of the model, one must experimentally manipulate the bump-position regulating ring neurons and the bump-width controlling ring neurons.

The neurons in the model are simulated using the leaky integrate-and-fire model and the synapses are conductance-based with the exponential dynamics. The interactions between EIP, PEI and PEN neurons are mediated through excitatory NMDA receptors. Ring neurons inhibit the EIP, PEI or PEN neurons though GABAergic receptors.

### The behavior model

The main goal of the model was to demonstrate whether the EB-PB circuits are capable of maintaining and updating the activity bump under the given experimental conditions so that an accurate orientation is available to the flies when they want to fixate on the landmarks or the cued directions. The EB-PB circuit model does not control when the flies decide to fixate and how they move their bodies. This steering control is likely to be carried out in the brain regions downstream to the EB-PB circuits (Stone et al., 2017) and are out of the scope of the present study. Therefore, we modelled the behavior of the fly using a simple mathematical model. By analyzing recorded fly behavior in the task, we found that the flies randomly switched between two behavioral states: forward movement and rotation, and the statistical characteristics matched those of the Markov-chain dynamics. Interestingly, we discovered that although the flies performed fixation during the stimulus and post-stimulus stages, the overall movement behavior in the two stages still matched the Markov-chain dynamics. This was because the flies in these two stages still randomly switched between the forward movement and rotation states. The fixation was produced by the occasional rotations that were made toward the landmarks. Based on the analyses we implemented a random movement protocol, which consists of two movement states, forward movement and rotation. The model switches between the two states randomly based on the Markov chain dynamics (see the section “The model parameters” below). In the model, a movement state lasted for a minimum length of one time step (300 ms). At the end of each time step, the fly switches to the other state or remains in the same state based on preset probabilities that are independent of the history. The probability of switching from forward movement to rotation is 0.40 and from rotation to forward movement is 0.60. These numbers were derived by analysing the distribution of bout length for each movement type in the behavioral tasks (see “The model parameters” section below).

During the forward movement state, the fly maintains its orientation without turning, while during the rotation state, the fly rotates in a randomly selected direction until the end of the state. The body rotation is accompanied by updating the activity bump in the EB-PB model through the unilateral activation of the PB.

### The simulation protocol

We simulated the second (stimulus) and the third (post-stimulus) stages of the behavioral task. In the simulation, the two stages were represented by landmark onset and offset, respectively. In the stimulus stage, the circuit model received visual input from the landmark and the signal about the movement state (forward or rotation) from the behavioral model described above. The visual input was modelled by sending spike input to the PB subregions which triggered an activity bump in the EB that corresponded to the landmark location. For the forward movement state, the P-ring neurons were activated to suppress the P circuit and the C circuit were allowed to operate in order to stabilize the activity bump in a fixed location. For the rotation state, the C-ring neurons were activated to suppressing the C circuit. At the same time, an input to the unilateral PB was applied in order to shift the activity bump in the P circuit. Input to the left side of the PB induced clockwise rotation of the activity bump, while input to the right side of the PB induced counterclockwise rotation. This input, presumably originated from the movement feedback, is used to model the observed counter-movement of the activity bump when the fly rotates its body. In the post-stimulus stage, the visual input was turned off, while the movement feedback remained. Approximately 200 trials were simulated for each experimental condition.

### The model parameters

The parameters of the EB-PB model mainly follow those used in a previous study (Su et al., 2017), but some parameters were modified in this study (Table 1). The parameters of the behavioral model were determined by the behavioral experiment described in this study. We analyzed the movement patterns of the flies during the first (pre-stimulus) stage and used them to determine the parameters associated with the forward movement and rotation state. By fitting an exponential curve to the distribution of the forward movement bouts, and the body rotation bouts, we determined the probabilities that a fly switch between the forward movement states and the rotation state. Based on the result, we constructed a Markov chain model of the random walk behavior of the fruit flies.

**Table 1.**
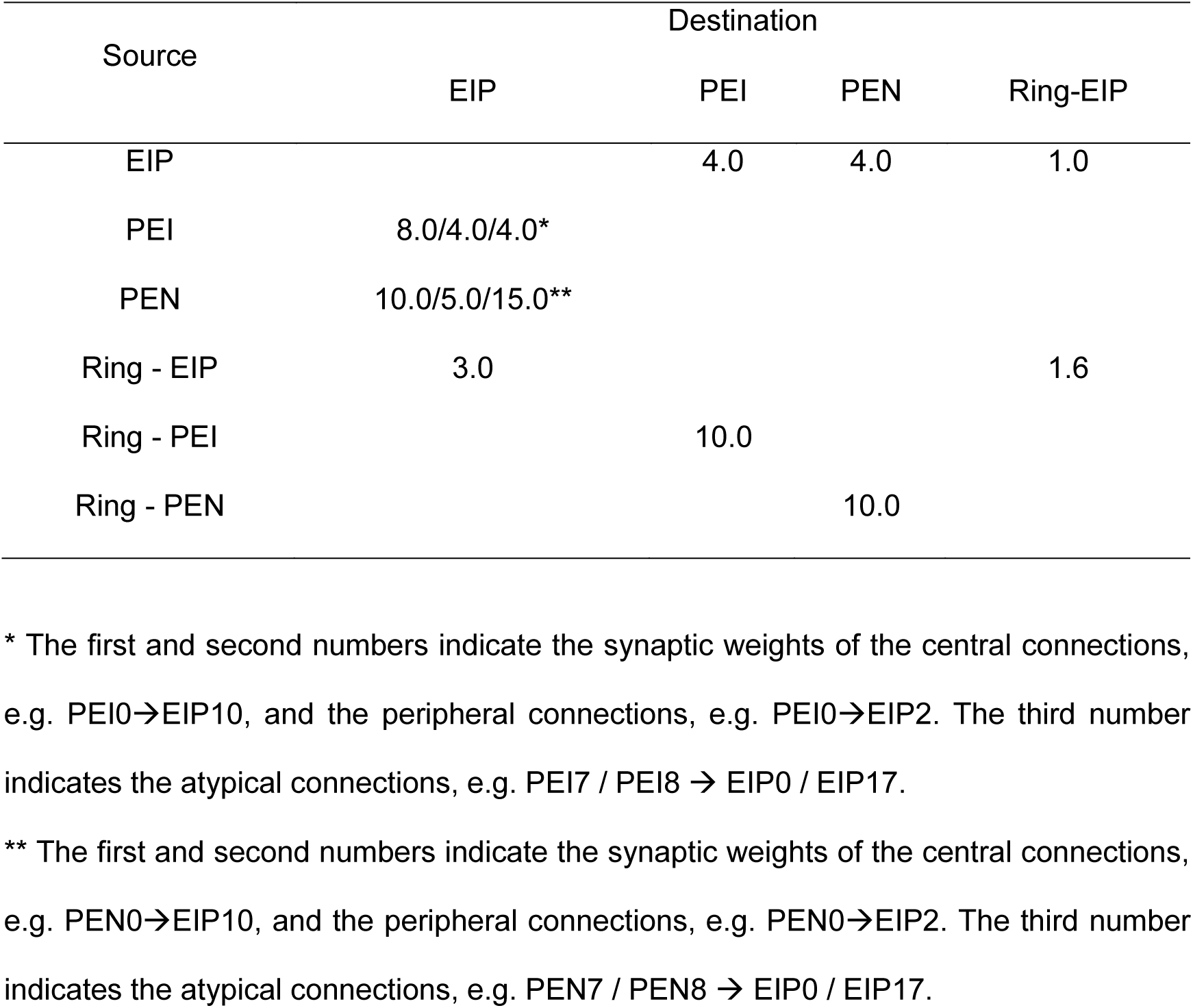
Synaptic weights between neuron types.

There were several parameters to be determined: The activation level of the EIP-ring and P-ring neurons in the ring-neuron suppression and photoactivation experiments. We left these as the free parameters and determined them by matching the resulting bump activity to the observed behavior in experiments. For the simulated flies with suppressed EIP-ring neurons, the input was -0.04 nA, while for the simulated flies with suppressed P-ring neurons, the input was reduced to -0.25 nA. For photoactviation of EIP-ring and P-ring neurons, the inputs were both 0.20 nA.

### Code accessibility

The major parameters are provided in Materials and Methods and Table 1. The full source code and parameter files are available at https://figshare.com/articles/online_resource/eNeuro_2020/13359041.

## Results

### Spatial orientation working memory

To investigate the mechanism of spatial orientation working memory in *Drosophila melanogaster*, we first developed a behavioral task based on Buridan’s paradigm (Götz, 1980; Strauss and Pichler, 1998; Yen et al., 2019) (see Materials and Methods) (Fig. 1A, B; Extended Data Fig. 1-2A). The task consisted of three stages. In the first (pre-stimulus) stage, in which no visual landmark was presented, the fruit flies walked randomly without any preferred orientation (Fig. 1D; Extended Data Fig. 1-2B) and spent most of the time walking along the edge of the arena (Extended Data Fig. 1-2C). In the second (stimulus) stage, in which two landmarks appeared at 0*°* and 180*°* on the screen, the flies exhibited significant visual fixation behavior by walking back and forth between the two landmarks and spent less time walking along the edge (Fig. 1E). This stage tested the ability of flies to orient themselves toward visual landmarks, and no memory was required. In the third (post-stimulus) stage, in which the landmarks disappeared, we found that the flies still maintained a behavior preference similar to that in the second stage, although with a weaker tendency of fixation (Fig. 1F). The ability of fixation in this stage indicated successful recall of the landmark directions with respect to the fly’s momentary heading, and is therefore an indication of spatial orientation working memory. To determine whether the strength of fixation depends on the stimulus (landmarks) exposure time, we conducted the task using different durations for the stimulus stage and calculated the fixation density *FD*, which indicates the percentage of time the flies spent in fixating on the landmark locations during a stage (see Materials and Methods) (Fig. 1G; Extended Data Fig. 1-4).

We observed that the fixation density in the third stage was larger than that in the first stage for all stimulus exposure times, indicating that fruit flies hold memory about the landmark directions. We can estimate the duration of memory in the third stage by multiplying *PI* (=*FD_3rd stage_* − *FD_1st stage_*) by the duration of the third stage. This number represents the change of the fixation behavior evoked by the earlier exposure to the stimulus in the second stage, and hence indicates the trace of memory. Taking the condition of a 120 s stimulus stage for example, *PI* was ∼0.20, the duration of the third stage was 120 s, and the estimated memory duration was ∼24 s. Two interesting properties were discovered in this test: first, *PI*, and hence the estimated memory duration was positively correlated with the stimulus duration (Fig. 1G), and second, the estimated memory duration is surprisingly long considering the stimulus duration was shorter than that used in previous studies (Neuser et al., 2008; Strauss and Pichler, 1998).

### Impairment of spatial orientation working memory by suppressing ring neurons

Next, we investigated how suppressing the EIP- or P-ring neurons may affect spatial orientation memory in the behavioral task. We used the same behavioral paradigm as described in the previous section with a 60 s stimulus stage and suppressed the ring neurons by hyperpolarizing the membrane (*;;UAS-Kir2.1*) (Hong et al., 2008) or by blocking synaptic transmission (*;;UAS-TNT*) (Hong et al., 2008) (see Materials and Methods). For EIP-ring neuron (Fig. 2A) suppression, we found that both methods led to similar behavioral changes. During the stimulus stage, the flies with suppressed EIP-ring neurons exhibited the same *PI* as did the wild-type flies (Fig. 2B). But during the post-stimulus stage, *PI* of the flies with suppressed EIP-ring neurons decreased significantly, showing that the flies did not fixate on the landmark locations after their offset (Fig. 2C).

**Figure 2.**
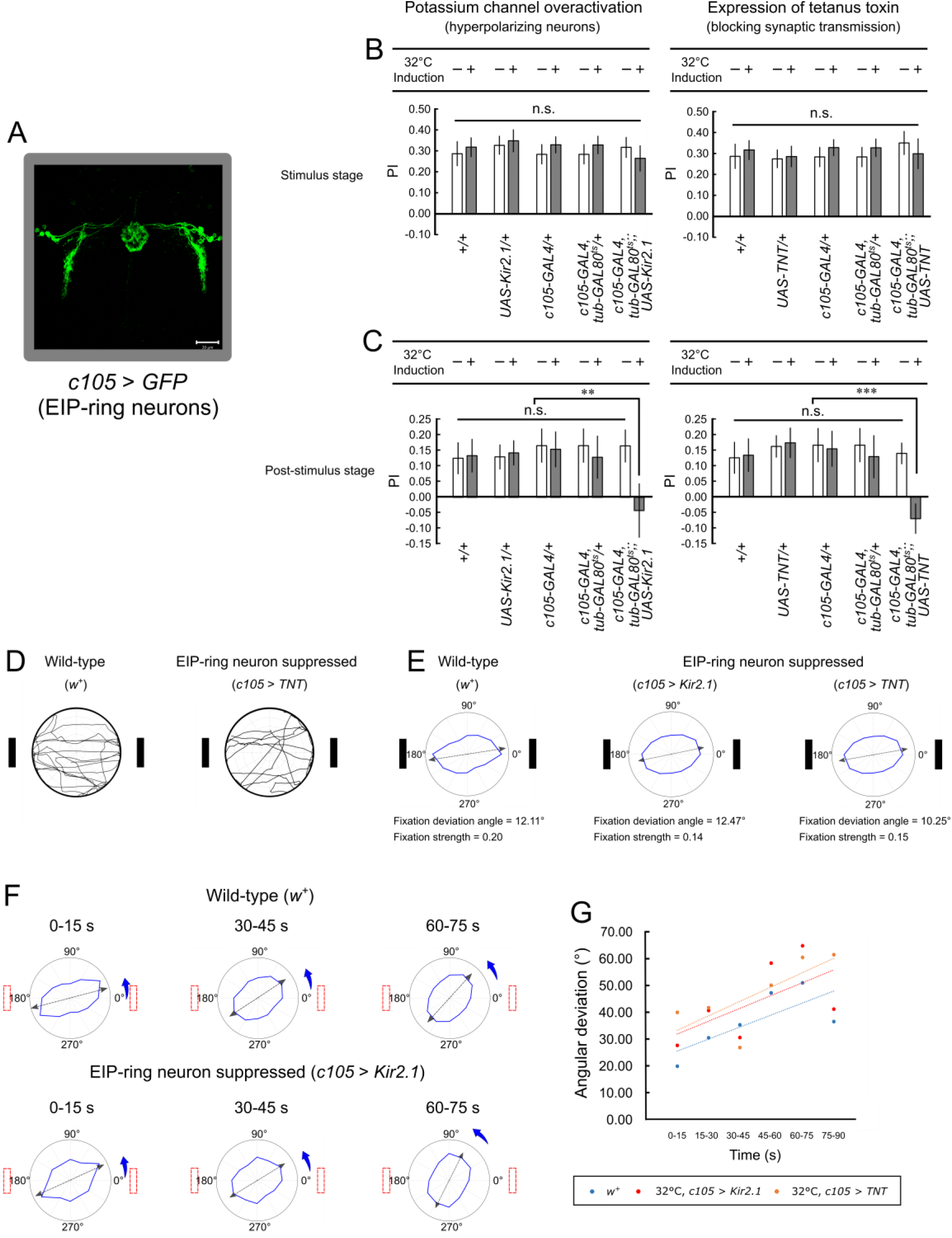
The fruit flies with suppressed EIP-ring neurons exhibited less-accurate orientation memory in the post-stimulus stage. ***A***, The *GAL4* expression domains in the EIP-ring neurons (*c105-GAL4;; UAS-GFP*). Eleven virtual slices are used in this confocal image. The thickness of each single slice is 0.90 *µm* and the thickness of all these 11 slices is 9.88 *µm*. ***B***, The performance index (*PI*) of wild-type flies and flies with suppressed EIP-ring neurons (left: 32°C, *c105-GAL4, tub-GAL80^ts^;; UAS-Kir2.1*. Right: 32°C, *c105-GAL4, tub-GAL80^ts^;; UAS-TNT*) in the stimulus stage. *, **, and *** indicate statistical significance with P-values of < 0.05, < 0.01, and < 0.001, respectively; n.s. = non-significant. ***C***, Same as in ***B***, but for the post-stimulus stage. ***D***, Example movement trajectories of a control fly (wild-type, *w^+^*) and a fly with suppressed EIP-ring neurons (32°C, *c105-GAL4, tub-GAL80^ts^;; UAS-TNT*) during the stimulus stage. ***E***, The radar plots of control flies and flies with suppressed EIP-ring neurons (both methods) in the stimulus stage indicate that the manipulation of EIP-ring neurons did not produce significant effect on fixation behavior ***F***, Significant effects on the fixation behavior during the post-stimulus stage can be observed for flies with suppressed EIP-ring neurons (32°C, *c105-GAL4, tub-GAL80^ts^;; UAS-TNT*) versus the control group (wild-type: *w^+^*). The radar plots are displayed for different time windows of the post-stimulus stage (Extended Data Fig. 2-1). ***G***, Fixation deviation angle as a function of time for the wild-type files and the flies with EIP-ring neuron suppression during the post-stimulus stage. Dashed lines indicate the linear regression of the color-matched data points. The deviation angle was significantly larger in flies with EIP-ring neuron suppression than in the wild-type flies.

The *PI* plots described above provide information regarding the frequency of movement toward the landmarks versus toward the perpendicular directions. However, it is also important to investigate the overall movement patterns of the flies in all directions. Inspection of the movement trajectories (Fig. 2D) and the radar plots (see Materials and Methods) of the wild-type flies and the flies with suppressed EIP-ring neurons (Fig. 2E) revealed that the suppression did not produce significant effect in the stimulus stage, indicating that the flies largely maintained their fixation behavior. However, the effect was significant during the post-stimulus stage. We discovered that while the wild-type flies exhibited a slight deviation from the actual landmark direction (Fig. 2F top row; Extended Data Fig. 2-1A), flies with suppressed EIP-ring neurons exhibited a profound change in the preferred movement direction during the post-stimulus stage (Fig. 2F bottom row; Extended Data Fig. 2-1B, C). This deviation increased with time (Table 2). Flies that underwent two different methods of EIP-ring neuron suppression exhibited similar deviation rates, which were markedly larger than that of the wild-type flies (Fig. 2G).

**Table 2.**
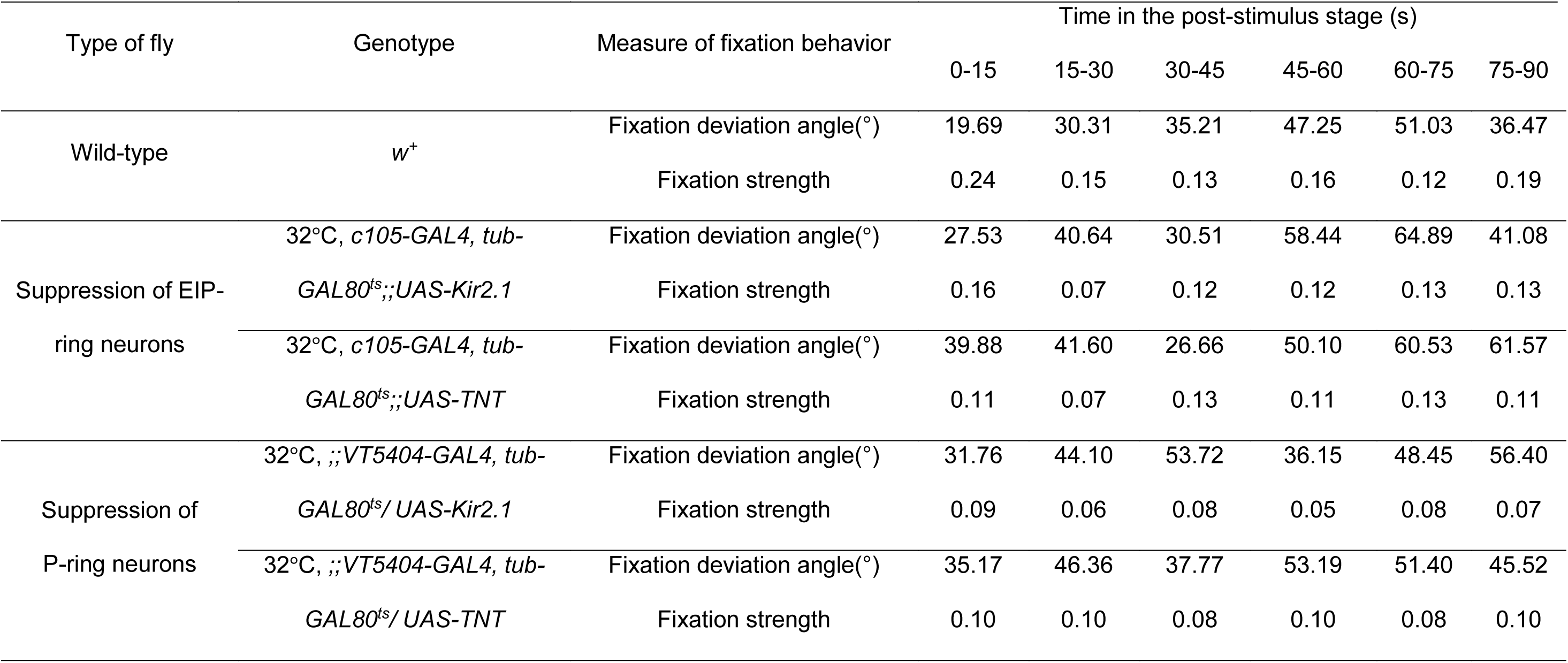
Fixation deviation and fixation strength of wild-type flies, flies with suppressed EIP-ring neurons, and flies with suppressed P-ring neurons in different time windows of the post-stimulus stage.

We next investigated the behavioral changes involving suppression of the P-ring neurons using the same UAS lines (*UAS-Kir2.1* and *UAS-TNT*) as used for the EIP-ring neurons (Fig. 3A). Whereas both methods of P-ring neuron suppression led to similar behavioral changes, they were distinct from those led by EIP-ring neuron suppression. With suppression of P-ring neurons, the *PI* of the flies decreased significantly in both the stimulus and post-stimulus stages (Fig. 3B, C). By inspecting the movement trajectories and the radar plots, we discovered that the flies with suppressed P-ring neurons moved randomly during the stimulus stage (Fig. 3D) and the post-stimulus stage (Fig. 3E; Extended Data Fig. 3-1).

**Figure 3.**
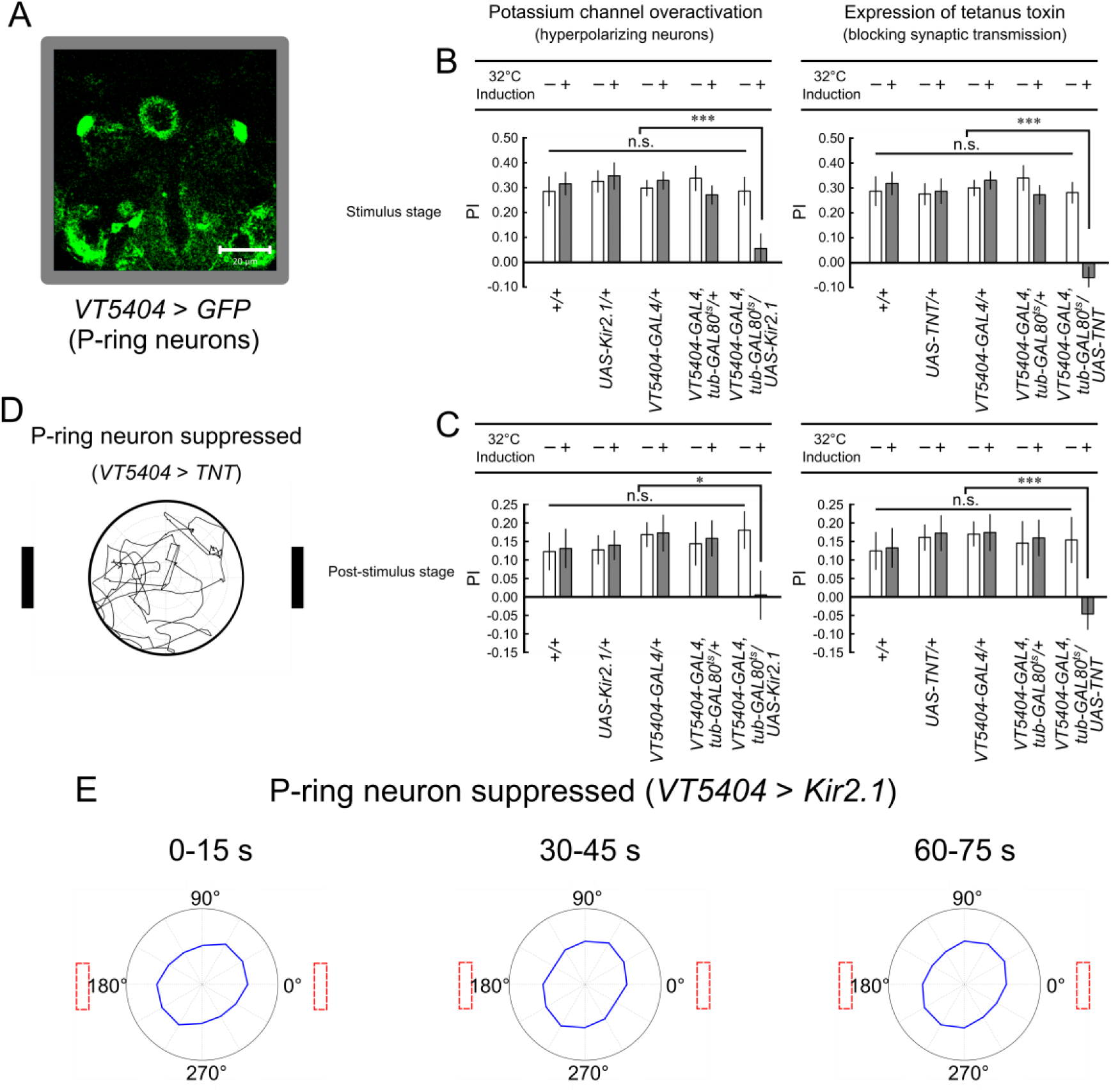
The fruit flies with suppressed P-ring neurons exhibited non-fixation behavior in both stimulus and post-stimulus stages. ***A***, *GAL4* expression domains in the P-ring neurons (*;;VT5404-GAL4/ UAS-GFP*). Nine virtual slices are used in this confocal image. The thickness of each single slice is 2.06 *µm* and the thickness of all these 9 slices is 18.52 *µm*. ***B***, The performance index (*PI*) of the control flies and flies with suppressed P-ring neurons (32*°*C, *;;VT5404-GAL4, tub-GAL80^ts^/ UAS-Kir2.1* & 32*°*C, *;;VT5404-GAL4, tub-GAL80^ts^/UAS-TNT*) in the stimulus stage. ***C***, Same as in ***B***, but for the post-stimulus stage. ***D***, Example movement trajectories of a fly with suppressed P-ring neurons (32*°*C, *;;VT5404-GAL4, tub-GAL80^ts^/ UAS-TNT*) in the stimulus stage. ***E***, Radar plots of flies with suppressed P-ring neurons (32*°*C, *;;VT5404-GAL4, tub-GAL80^ts^/ UAS-Kir2.1*) in different time windows of the post-stimulus stage (Extended Data Fig. 3-1). *, **, and *** indicate statistical significance with P-values of < 0.05, < 0.01, and < 0.001, respectively; n.s. = non-significant.

One may argue that the loss of fixation behavior in flies with suppressed P-ring neurons was not the result of impairment in spatial orientation but was due to deficiencies in other functions such as vision or locomotion. We first tested the vision for the flies with suppressed P-ring neurons by conducting the laser escaping task (see Materials and Methods). We measured the escape rate for the approaching laser beam and found no significant difference between the wild-type flies (71.88%) (Movie 3), and the flies with suppressed P-ring neurons (80.95%) (Movie 4), while the escape rate of the fruit flies with deficient photosensors (genotype: *;;ninaE*) (Movie 1) was 0%, suggesting that P-ring neuron suppression did not impair vision.

To investigate whether the motor function was normal in flies with suppressed P-ring neurons, we measured the activity level (percentage of movement bouts in a given period) and speed (mean movement speed in the movement bouts) in the pre-stimulus stage. We did not observe significant differences between the wild-type flies and the flies with suppressed P-ring neurons (Table 3). The analysis of speed and activity level may not reflect subtle movement impairments such as steering difficulty. Therefore, we performed detailed analysis on the responses of flies in the laser escaping task described above. The flies exhibited three major types of escape maneuvers in the task: detour, turnback and acceleration. We found that the percentages of each type of escape maneuver are comparable between the wild-type flies and flies with suppressed P-ring neurons (Table 4). For comparison, we also tested flies with steering difficulty by amputating the unilateral foreleg of wild-type flies (Isakov et al., 2016). The flies had tremendous difficulty to escape the approaching laser beam and the escape rate was only 10.37% (Movie 2).

**Table 3.**
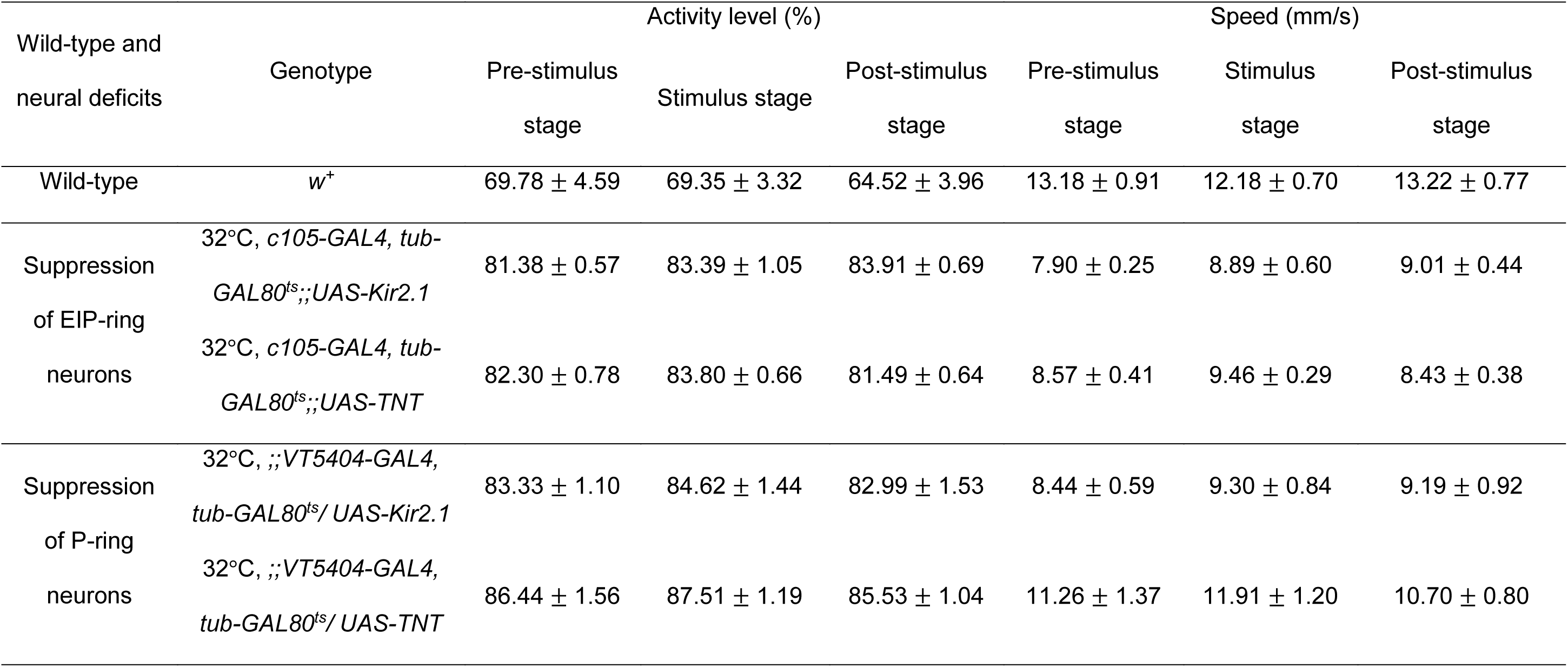
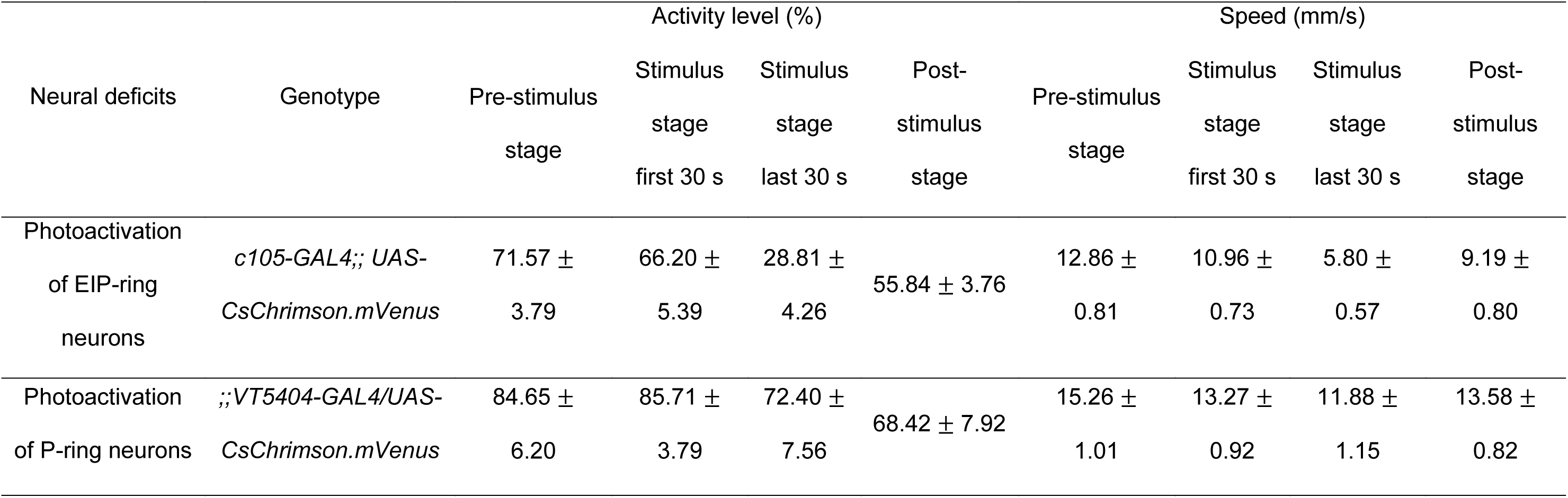
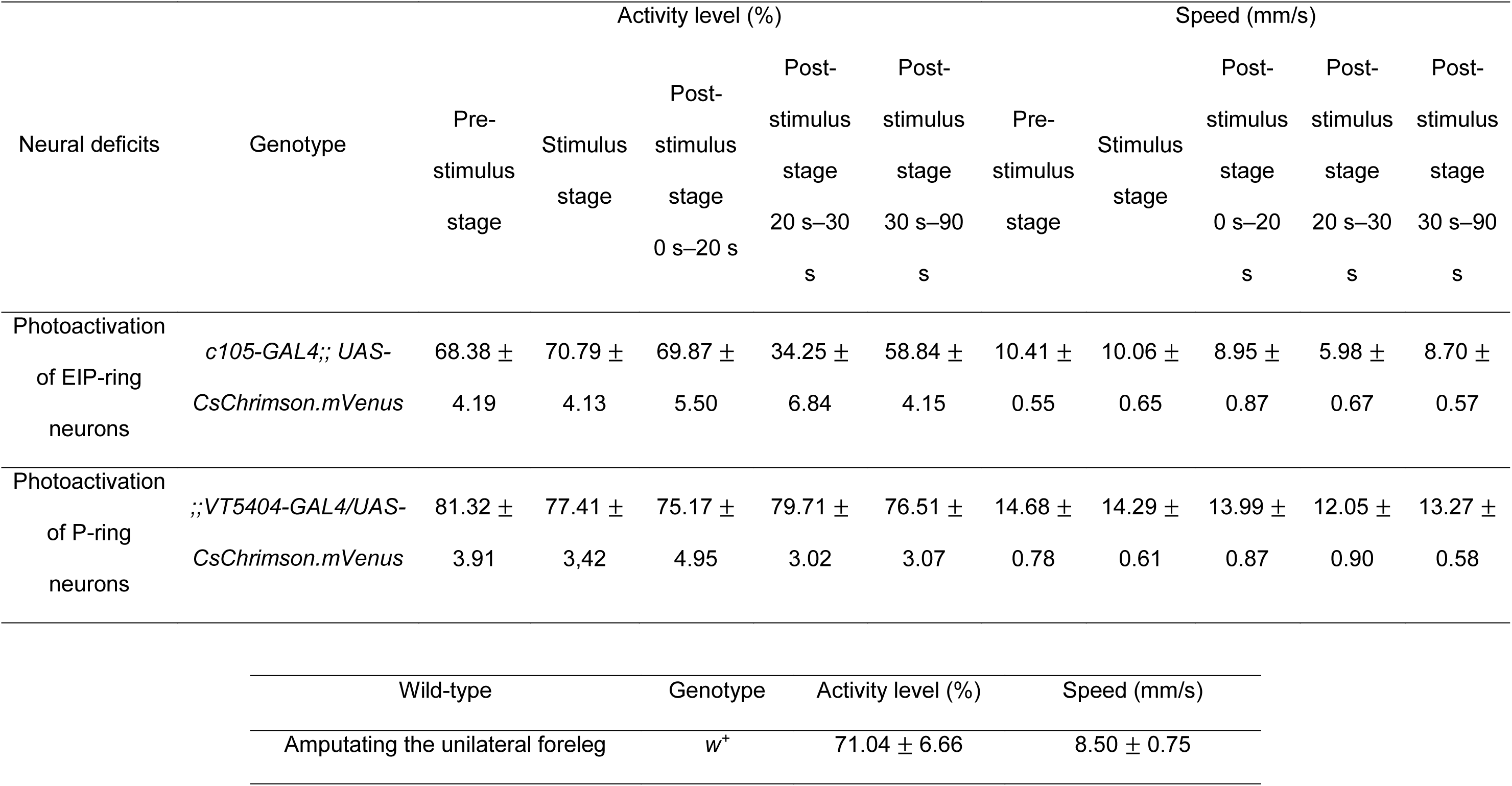
Activity level and movement speed of wild-type flies, flies with suppressed EIP-ring or P-ring neurons and flies with photoactivated EIP-ring or P-ring neurons.

**Table 4.**
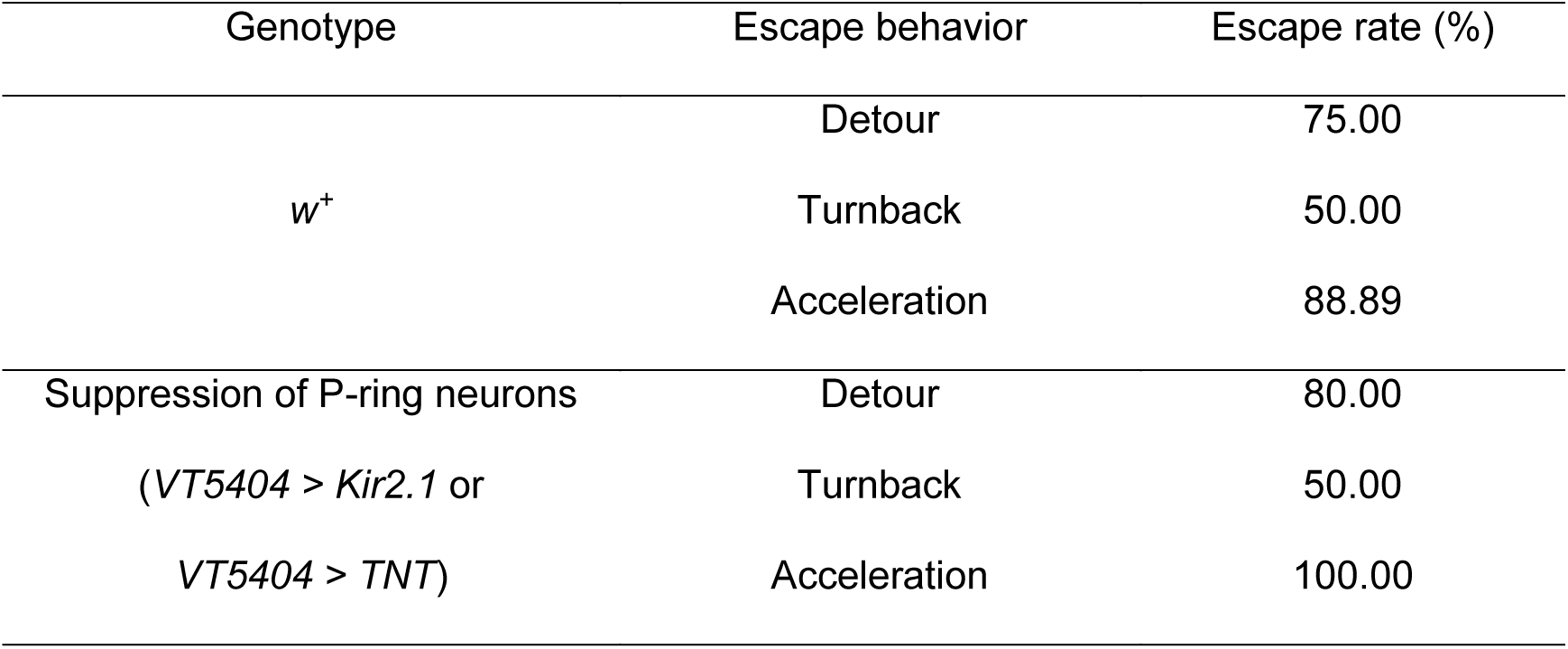
The percentages of each type of escape maneuver in the laser escaping task for the wild-type flies and flies with suppressed P-ring neurons.

### Impairment of spatial orientation working memory by activating ring neurons

We have inspected the effect of the ring neuron suppression, which presumably overactivated the downstream circuits. But how about suppression of these downstream circuits? In particular, if we transiently suppress the circuits, would orientation memory interrupt? To this end, we activated the EIP-ring or P-ring neurons by optogenetics using two temporal protocols (see Materials and Methods) (Fig. 4; Extended Data Fig. 4-1 and 4-2). For EIP-ring neuron photoactivation during the last 30 s of the stimulus stage, the movement speed and activity level of the flies significantly reduced during this period (Table 3; Movie 5), but the fixation behavior in the post-stimulus stage did not exhibit significant difference with that of the wild-type flies (Fig. 4A). The result indicated that orientation working memory is not interfered by photoactivation of the EIP-ring neurons during the stimulus stage. When the photoactivation was applied midway during the post-stimulus stage (20-30 s), the flies lost the fixation behavior after the photoactivation (Fig. 4B). For P-ring neuron photoactivation during the last 30 s of the stimulus stage, the speed and activity level of the flies were close to that of the wild-type flies (Table 3; Movie 6), and the fixation behavior during the activation period remained intact but slightly weakened during the post-stimulus stage (Fig. 4C). However, photoactivating the P-ring neurons during the 20-30 s of post-stimulus stage abolished the subsequent fixation behavior (Fig. 4D), an effect that is identical to that of the EIP-ring activation during the same period (Fig. 4B, E).

**Figure 4.**
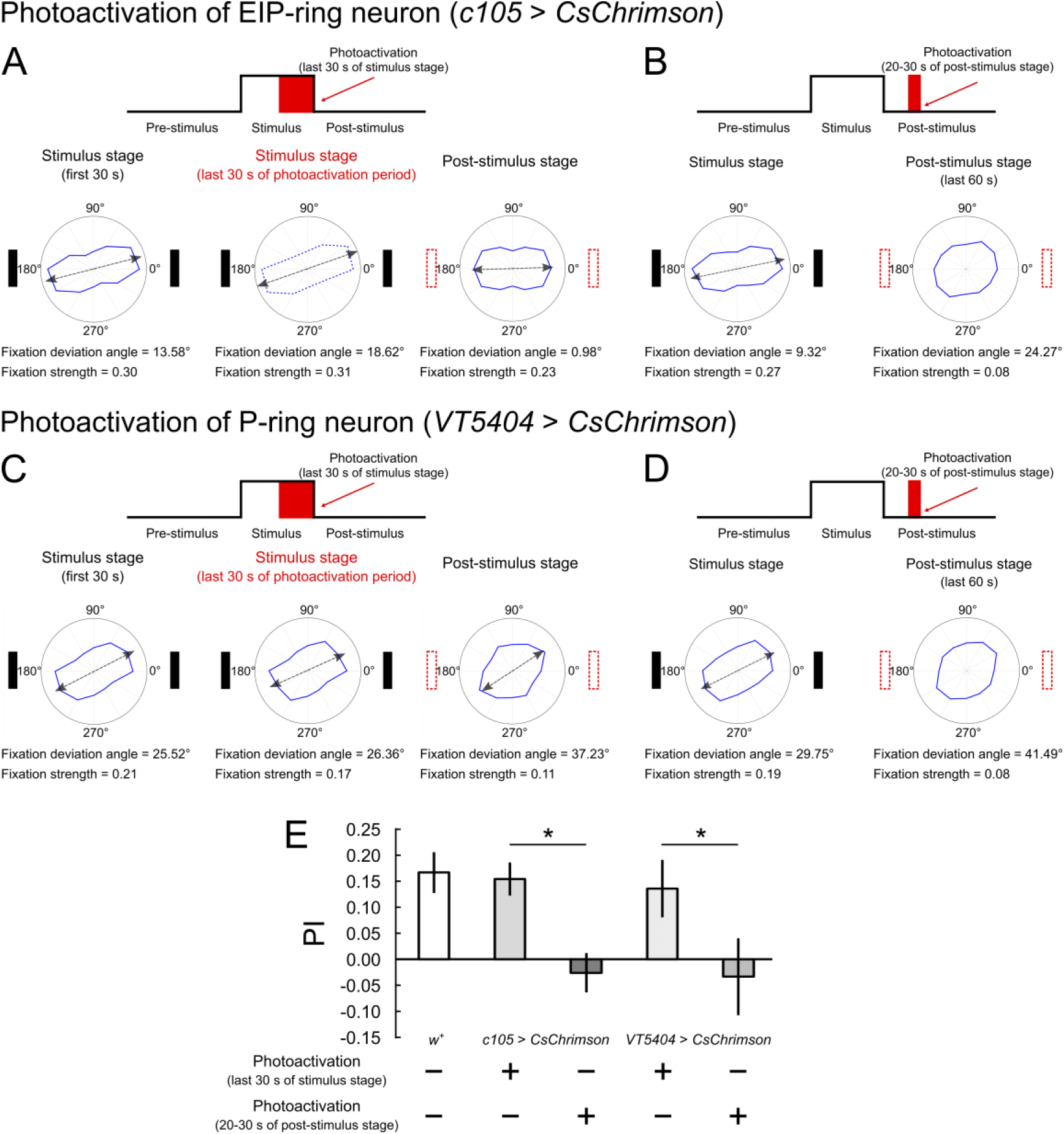
Impact of transient photoactivation of the EIP-ring or P-ring neurons on the spatial orientation working memory. ***A***, Photoactivation of the EIP-ring neurons (*c105-GAL4;;UAS-CsChrimson.mVenus*) during the last 30 s of the stimulus stage. Top: the stimulation protocol. Bottom: The radar plots for the first 30 s of the stimulus stage (left bottom), the last 30 s of the stimulus stage where the photoactivation was applied (middle bottom) and the post-stimulus stage (right bottom) indicate that the orientation memory was intact. Note that the flies performed very little movement during the period of the photoactivation (Table 4). ***B***, Same as in ***A***, but with photoactivation during 20-30 s of the post-stimulus stage. The radar plots are shown for the stimulus stage and the last 60 s of the post-stimulus stage. The fixation behavior was abolished after the photoactivation. ***C-D***, Same as in ***A-B***, but with photoactivation of the P-ring neurons (*;;VT5404-GAL4/ UAS-CsChrimson.mVenus*). The photoactivation during the post-stimulus stage also abolished the fixation behavior. ***E***, The *PI* of wild-type flies (*w^+^*) and flies in each photoactivation condition. Photoactivation in the stimulus stage caused minor or no impact on the orientation memory while photoactivation in the post-stimulus stage abolished the memory. Performance of control groups (no photoactivation) are shown in Extended Data Figure 4-1, 4-2 and 4-3.

### Hypothesis of the underlying neural mechanisms

To summarize the results of the behavioral and neural functional tests, we plot schematics that illustrate the observed movement patterns of wild-type flies and flies with EIP-ring or P-ring neuron manipulation in all three stages (Extended Data Fig. 5-1). The wild-type flies exhibited strong fixation behavior toward the landmark directions during the stimulus stage. The fixation behavior was still strong but with a slight deviation in the fixation direction during the post-stimulus stage, indicating the presence of working memory of the landmark direction (Extended Data Fig. 5-1A). Flies with photoactivation of EIP-ring or P-ring neurons during the stimulus stage performed similarly (Extended Data Fig. 5-1A). Flies with suppressed EIP-ring neurons also exhibited a strong preference during the stimulus stage (Extended Data Fig. 5-1B). However, comparing to the wild-type flies, the deviation increased progressively during the post-stimulus stage, suggesting that the flies maintained the memory of the existence of the landmarks but with incorrect memory of their orientation. By contrast, the flies with suppressed P-ring neurons did not exhibit fixation behavior in both stages, indicating that these flies lost spatial orientation regardless the visibility of the landmarks (Extended Data Fig. 5-1C). Finally, short photoactivation during the post-stimulus stage of either EIP-ring or P-ring neurons both led to abolished fixation behavior after the activation period (Extended Data Fig. 5-1D).

The behavioral results can be explained by the neural mechanisms proposed in the EB-PB model (Su et al., 2017) (Fig. 5). See Materials & Methods for a brief description of the model mechanisms. To visualize the effects of neural manipulations on the circuits, we simplify the model diagram into an abstract three-ring representation and each ring is modulated by one of the three ring neuron types (C-ring, P-ring and EIP-ring) (Fig. 5A, B).

**Figure 5.**
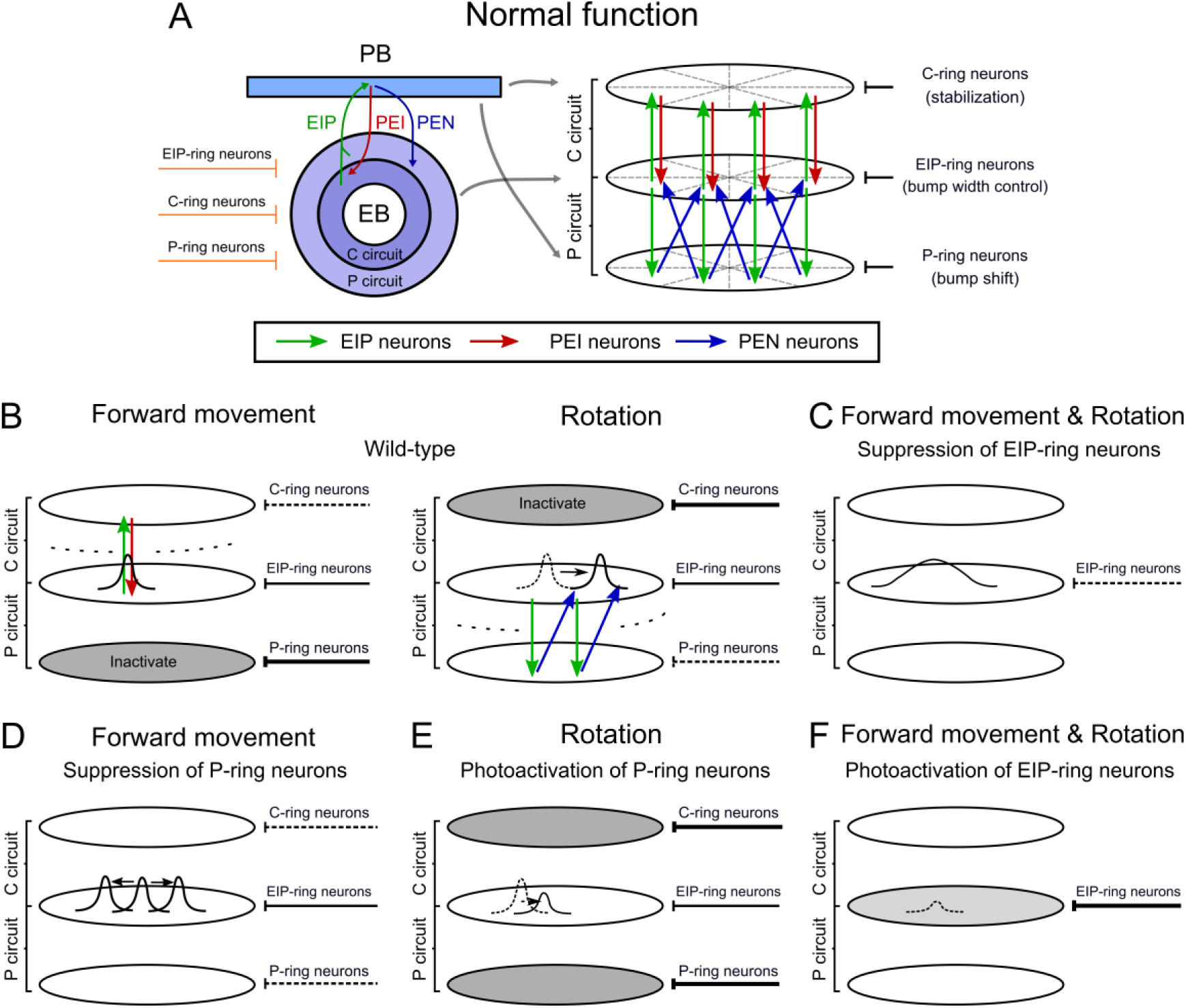
Schematics of the neural activity underlying the behavioral effects of the EIP-ring or P-ring manipulation based on the EB-PB circuit model (Su et al., 2017). ***A***, The EB-PB circuit model consists of a C circuit (EIP & PEI neurons) and a P circuit (EIP & PEN neurons). The former stabilizes the localized neural activity (the bump) that represents the direction of the landmark and the latter updates the memory by shifting the bump when the body rotates. ***B***, When a fly moves forward, the C circuit is activated and the P circuit is inhibited by the GABAergic P-ring neurons (left). On the other hand, when the fly rotates, the C circuit is inhibited by the GABAergic C-ring neurons and the P circuit is partially activated (right). The GABAergic EIP-ring neurons are always activated in order to maintain the width of the bump through the global inhibition. ***C***, When the EIP-ring neurons are suppressed, the activity bump becomes wider and drifts more, which leads to an inaccurate orientation memory. ***D***, When the P-ring neurons are suppressed, the P circuit becomes fully activated even during forward movement. The overactivation leads to a destabilized bump that diffuses throughout the ring and the fly loses its orientation. ***E***, The photoactivation of the P-ring neurons suppresses the P circuit, which loses the activity bump during rotation. ***F***, The photoactivation of EIP-ring neurons produces overly powerful global inhibition that suppresses the activity bump during the post-stimulus stage when no landmark input is available to support the bump. Schematics of the observed behavior changes is shown in Extended Data Figure 5-1.

Based on the model, if the EIP-ring neurons are suppressed, the reduction of inhibition leads to a broadened bump with jittered position (Fig. 5C), which makes the orientation memory less accurate (Extended Data Fig. 5-1B). When the P-ring neurons are suppressed, both clockwise and counterclockwise part of the P circuit becomes activated, shifting the bump toward both directions. In consequence, the neural activity spreads through the entire ring (Fig. 5D) and the fly loses its orientation completely (Extended Data Fig. 5-1C). If we photoactivate P-ring neurons during the post-stimulus stage, then the P circuit is suppressed even during body rotation when the C circuit is also inhibited. Therefore, the bump cannot be sustained because of the lack of recurrent excitation from the C and P circuit (Fig. 5E). As a consequence, the fly loses its orientation memory (Extended Data Fig. 5-1D). Finally, if the photoactivation is performed on the EIP-ring neurons during the post-stimulus stage, it would diminish the activity bump due to the excessive inhibition (Fig. 5F). Therefore, the fly loses its orientation memory in the third stage (Extended Data Fig. 5-1D). However, if the photoactivation is performed during the second stage in which the landmarks are visible, the bump could still be maintained by the input corresponding to the visual cue (Extended Data Fig. 5-1A).

### The computational model of spatial orientation working memory

We have illustrated in Figure 5 how interrupted functionality of the C or P circuits affected the spatial orientation working memory based on our basic understanding of the dynamics of the EB-PB model (Su et al., 2017). Next, we further demonstrated such neural mechanisms by performing the model simulations based on the same experimental protocols used in the present study. The model consists of the EB-PB neural circuit model and a Markov-chain behavioral model (see Materials and Methods; Extended Data Fig. 6-1).

We first showed that the model with default settings (corresponding to wild-type flies) was able to maintain activity bump in both stimulus and post-stimulus stages as expected (Fig. 6A). The model with suppressed EIP-ring neurons still exhibited a clear activity bump (Fig. 6B) in both stimulus and post-stimulus stages while P-ring neuron suppression lost the activity bump completely in both stages (Fig. 6C). However, compared to the wild-type, although the bump was maintained in the condition of EIP-ring neuron suppression, the bump width (*FWHM* = 2.04 ± 0.04 rad) is larger than that of the simulated wide-type flies (*FWHM* = 1.86 ± 0.05 rad). Similar to the real flies, the simulated wild-type files exhibited progressively increasing deviation between the bump position and the head direction during the post-stimulus stage (Fig. 6D). Moreover, the bump with EIP-ring neuron suppression drifted more during the post-stimulus stage and the mean deviation between the true heading and the activity bump location was larger than the simulated wild-type fly (Fig. 6E). The simulated result is consistent with what are pictured in Figures 5B-5D, and explained the observations shown in Figures 2 and 3. Flies with suppressed EIP-ring neurons still performed robust fixation activity due to the existence of the activity bump but with less accurate fixation direction due to the larger deviation between the true heading and the bump position. On the other hand, flies with suppressed P-ring neurons did not fixate due to the loss of the activity bump.

**Figure 6.**
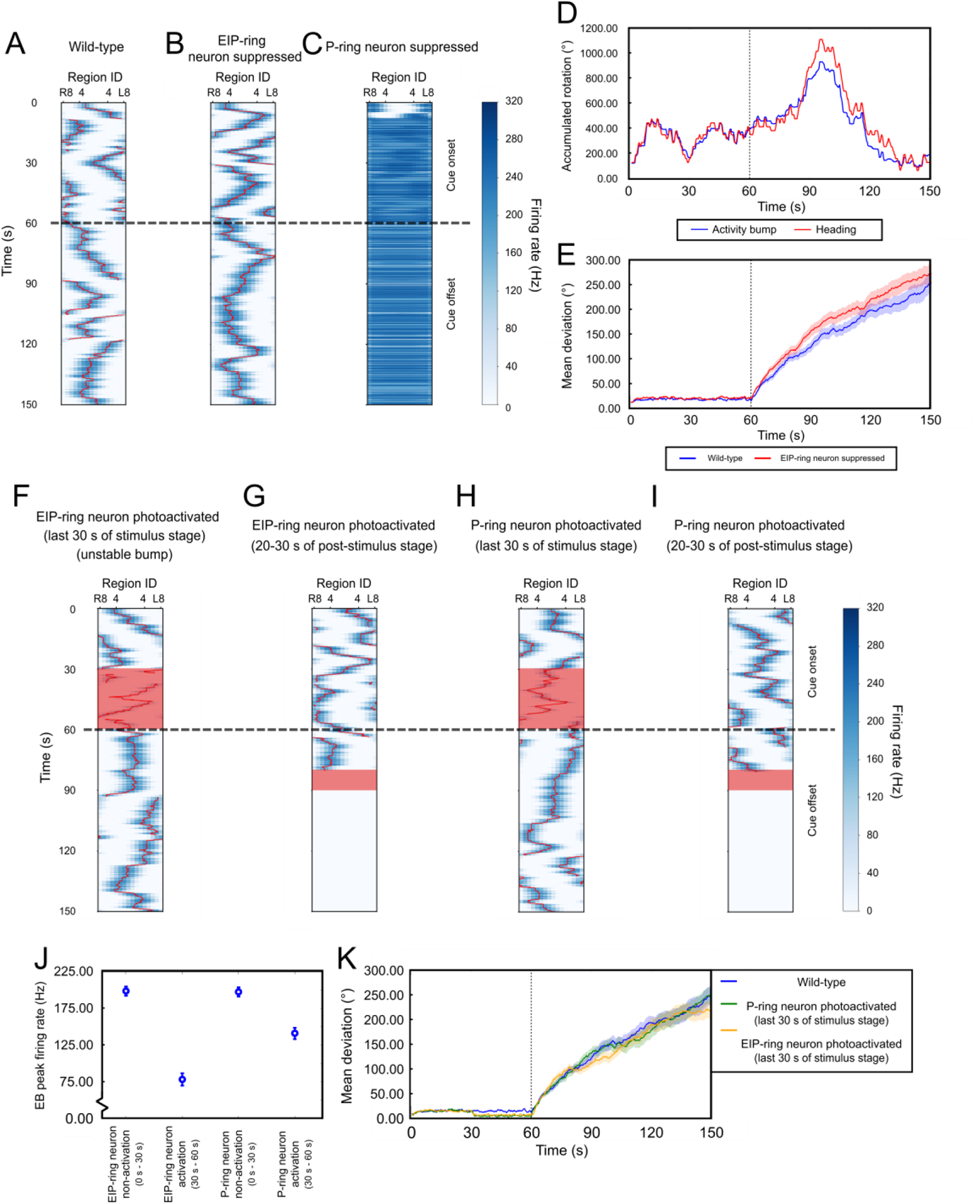
The computer simulations of the EB-PB model (Su et al., 2017) demonstrated the neural mechanisms illustrated in Figure 5, and highlighted the roles of the EIP- and P-ring neurons in the spatial orientation working memory. ***A***, The neural activity of EIP neurons in all EB regions as a function of time indicates the existence of activity bump (dark blue) in a simulated wild-type fly. The landmark was on between t = 0 s and t = 60 s (dashed line). ***B-C***, Same as in ***A***, but with simulated EIP-ring (***B***) and P-ring neurons (***C***) suppression. The P-ring neuron suppression led to unstable and widespread neural activity. ***D***, The accumulated angles of the true heading and the activity bump of in one example trial for a simulated wild-type fly. ***E***, The trial-averaged angular deviation between the head direction and the bump for simulated wild-type flies (blue) and simulated flies with suppressed EIP-ring neurons (red). Shades represent the standard error of the mean. ***F-G***, Same as in ***A***, but with simulated photoactivation (red regions) of the EIP-ring neurons. After the photoactivation in the last 30 s of the stimulus stage (***F***), the activity bump persisted in all trials. The photoactivation in the post-stimulus stage abolished the activity bump (***G***). ***H-I***, Same as in ***F-G*** but with the photoactivation of the P-ring neurons in the last 30 s of the stimulus stage (***H***) or during the post-stimulus stage (***I***). ***J***, EB peak firing rate with no-photoactivation and photoactivation of EIP-ring or P-ring neurons in the stimulus stage. ***K***, The trial-averaged angular deviation between the heading and the bump for simulated wild-type flies (blue), P-ring neuron photoactivation (green) and EIP-ring photoactivation (red). Shades represent the standard error of the mean. The behavior model used to drive the EP-PB circuit model is shown in Extended Data Figure 6-1.

Next, we simulated the photoactivation experiments in the model. The EIP-ring neuron activation during the last 30 s of the stimulus did not affect the activity bump during the post-stimulus stage (Fig. 6F), while photoactivation during the post-stimulus stage completely abolished the bump (Fig. 6G). This is consistent with the picture depicted in Figure 5F and the experimental observation (Fig. 4B). Interestingly, P-ring neuron activation in the simulations produced similar effects (Fig. 6H, I) but with two subtle differences. First, when EIP-ring or P-ring neurons were activated during the last 30 s of the stimulus stage, the strength of the bump, as measured by the peak firing rate of the bump, was both reduced. But the reduction is much more significant for the activation of EIP-ring neurons than for the P-ring neurons (Fig. 6J). We do not know how this difference may be reflected at the behavioral level, but we did observe that the activity level during the period of photoactivation was lower in the EIP-ring neuron activation condition than in the P-ring neuron activation condition (Table 2, bottom two rows) in experiments. However, we stress that the causal relationship between the bump strength and the activity level is just a speculation and is not a prediction of the EB-PB model. Second, with P-ring neuron photoactivation during the second stage, we found that the bump could be lost in a small percentage (24.2%) of trials. This is due to the instability occurring when the P circuit recovered from the inhibition at the end of the photoactivation. The instability might result from the model implementation or the choice of model parameters, and might not reflect an actual neuronal phenomenon. Finally, for either EIP-ring or P-ring neuron activation, if the bump was maintained during the post-stimulus stage, the deviation between the true heading and the bump location is comparable to that in the simulated wild-type flies (Fig. 6K). This is consistent with the observation that the flies in these two conditions performed as good as the wild-type flies during the post-stimulus stage (Fig. 4E).

## Discussion

In the present study, we hypothesized that the C circuit and P circuit in the ellipsoid body circuits stabilize and update the orientation-encoding activity bump and they are regulated by corresponding GABAergic ring neurons. We tested our hypothesis by manipulating two types of GABAergic ring neurons in a spatial orientation working memory task with free moving fruit flies, and discovered manipulating each ring neuron type led to different behavioral abnormality. By performing computer simulations on a previously proposed EB-PB neural circuit model, we were able to explain the results of the experiments and provided a picture of the neural circuit mechanism underlying spatial orientation working memory: the orientation-encoding bump is maintained through two alternately activated neural processes: one that stabilizes the position of the activity bump and one that updates the position of the bump. The former is activated when a fruit fly maintains a steady head direction and the latter is activated when the fly rotates its body. The control of this process is performed through specific GABAergic ring neurons. Therefore, overactivating or suppressing the ring neurons disrupts the alternation of the two processes and leads to incorrect or even loss of orientation memory.

There are a few more interesting discoveries worth discussing. Performing fixation toward previous landmark directions requires two things to be remembered: the earlier event of the landmark presentation (what) and the directions of the landmarks (where). Flies that fail to remember the former would not exhibit the fixation behavior at all, while flies that forget the latter would still perform the fixation but toward incorrect directions. Our discoveries of strong fixation but with large deviation from the true directions of the landmarks for flies with EIP-ring neuron suppression during the third stage (Fig. 2F, G) may imply the segregation of the neural mechanisms of orientation memory regarding the “where” and “what” of a landmark.

One interesting finding of the present study is a long duration of spatial orientation working memory during the post-stimulus stage. Previous studies reported the occurrence of post-stimulus fixation behavior that lasted only for a few seconds immediately following the offset of the landmarks (Neuser et al., 2008). Indeed, we observed that the flies tended to stop their movement a few seconds after the sudden disappearance of the landmarks in the third stage. But they usually resumed the movement in a few seconds. This might be the reason why earlier studies only claimed a few seconds of fixation if their analyses did not include the resumed movement. Further studies are needed to investigate this issue.

A couple issues regarding the choice of molecular tools should be discussed. In the present study we used *tub-GAL80^ts^* in combination with *UAS-Kir2.1* or *UAS-TNT* to suppress targeted ring neurons. The method involves raising the temperature one day prior to the behavioral experiments and therefore taking effects on a much longer time scale than using optogenetic tools. This long-term suppression may induce other effects at the cellular or circuit levels, which are beyond what our model can simulate. Further study may be required to carefully examine the long-term effects. An ideal solution is to transiently suppress the ring neurons using optogenetic tools such as *UAS-NpHR* (peak sensitivity wavelength ≈ 589 nm) (Deisseroth, 2010) or *GtACRs* (peak sensitivity wavelength ≈ 527 nm for *GtACR1* and ≈457 nm for *GtACR2*) (Mauss et al., 2017). However, the wavelength of the required activation light is within the visible range of the fruit flies. Our preliminary tests on *UAS-NpHR* showed that the onset of the activation light seriously disrupted the fixation pattern of wild-type flies. A new optogenetic tool or a carefully re-designed optical system is required in order to transiently suppress targeted ring neurons while not interfering the visual experiments. The second issue is related to the *GAL4* lines. In the present study we only used two most specific lines, *c105-GAL4* and *VT5404-GAL4*, to target the EIP- and P-ring neurons, respectively. As listed in the Methods section, there are several other less specific *GAL4* lines available for the two type of neurons. It is necessary to conduct the same experiments using these overlapping lines to further confirm the results presented in this study.

Several other important questions remain to be addressed. Previous studies showed that EB does not maintain a fixed retinotopic map and a bump can start from a random location in the beginning of a trial (Seelig and Jayaraman, 2015). For the sake of modelling simplicity, we did not model the random starting point feature in our model. But this feature is easy to implement and does not affect the conclusion of this study. We simply need to apply a global excitation to the entire EB to reset the system. The excitation will induce strong competition between the EIP neurons and a new bump will start at a random location through the winner-take-all dynamics. Following this issue, random regeneration of an activity bump also needs to be discussed. Our model showed that photoactivation of either EIP-ring or P-ring neurons during the post-stimulus stage permanently abolished the activity bump. However, based on the observation of spontaneous generation of activity bump in other studies (Seelig and Jayaraman, 2015), the bump is likely to be regenerated at a random location after the offset of photoactivation. The regeneration can be easily implemented in our model using the same mechanism described above. Since the regenerated bump starts from a random location, the fruit flies lose the reference to the landmark locations. So adding a spontaneous bump or not both lead to the same conclusion: the files fail to fixate on the previous landmark locations. Although not affecting the conclusion of the present study, the spontaneous bump feature may be crucial in future studies that involve modelling of the steering mechanism.

Another issue is that a couple experimental and modelling studies suggested that ring neurons provides the mechanisms underlying flexible retinotopic mapping in EPG (or EIP) neurons rather than the simple suppression/activation mechanism as hypothesized in the present study (Fisher et al., 2019; Kim et al., 2019). However, the ring neurons (R2 & R4d) tested in one study (Fisher et al., 2019) are of different types from what we tested (R1 and R6) here. Further experiments that measure the activities of R2 & R4d using the setups described in Fisher et al. 2019 or Kim et al. 2019 are required to clarify this issue.

It is important to compare and discuss differences between computational models of the central complex in terms of the functions investigated in the present study. However, most models focused on different aspects of the compass circuit functions (see Introduction). Among these models the one proposed in Kakaria and de Bivort, 2017 (and was also used in Pisokas et al., 2020) can be compared directly to one used in Su et al., 2017 and the present study. Both models used biologically realistic spiking neurons and the underlying circuit mechanisms are comparable. The major difference is that Kakaria and de Bivort model proposes the PB intrinsic neurons as the main source of inhibition that regulates the attractor dynamics, while in our model this function is carried out by the EIP ring neurons with two additional ring neuron types (C and P-ring neurons) modulating different subcircuits of the system. An in-depth model comparison and experimental manipulation of PB intrinsic neurons and ring neurons under the present behavioral task may be able to clarify this issue.

A final issue is related to the function of the activity bump which is commonly thought to represent the fly’s sense of orientation in a manner similar to that of the head-direction system found in rodents (Muller et al., 1996). However, as aforementioned “what” and “where” mechanisms, performing the fixation behavior as an indication of orientation working memory may require several serial or parallel neural components beyond EB and PB. For example, how is this innate fixation behavior initiated (motivation)? When a fly stops fixating, it is not clear whether the fly forgets the landmark directions or simply enters a different behavior state (but still remembers the landmark directions). It also remains unclear whether the memory is stored in another neural circuit and the EB merely provides a reference frame for orientation, or whether the activity bump in the EB represents the actual memory of the landmarks. The current experimental setup is not able to address this issue. A novel task that can disassociate these two components is required for further investigation.

We conclude the present study as follows. First, the experiment indicated that long-term suppression of EIP-ring neurons reduced the accuracy of orientation working memory (fixation with an increased deviation angle), whereas long-term suppression of P-ring neurons abolished the memory completely (no fixation). Similarly, transiently activating either ring neuron types in the absence of landmark immediately abolished the memory. Second, the experimental observation can be explained by the EB-PB neural circuit model in which the EIP-ring neurons are responsible for controlling the width of the bump and the P-ring neurons are responsible for shifting (updating) the position of the bump. Third, put the experiment and the theory together, the present study suggests that coordinated activation of the two ring neuron types which control the downstream EB-PB subcircuits is crucial for spatial orientation working memory.

## Author Contributions

RH, CLC, HYC and CCL designed research; RH and HPH performed research; HHY and WTK contributed unpublished reagents/ analytic tools; RH and HPH analysed data; RH and CCL wrote the paper.

## Acknowledgements

We thank Dr. Ann-Shyn Chiang’s lab for providing several transgenic flies and thank him for his helpful discussion. We thank FlyLight Project Team at Janelia Research Campus for the use of the images. We also thank Jia-Ying Chien and Tzu-Yu Tseng for acquiring confocal images, and Tzu-Min Wei for the help in the behavioral experiments, and Ta-Shun Su for programming.

## Extended Data Figures

**Extended Data Figure 1-1.**
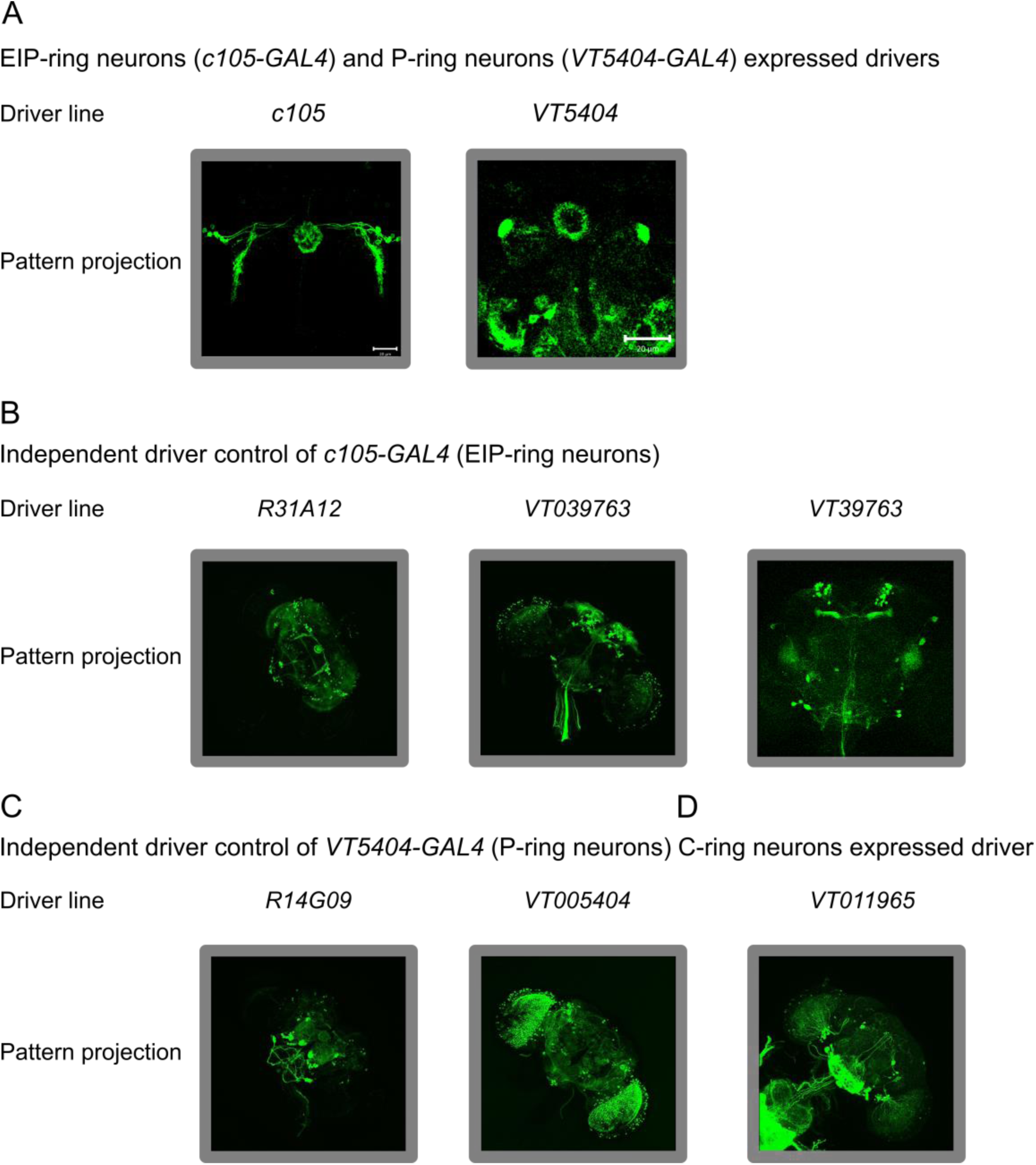
The fluorescent images of the *GAL4* drivers that are expressed in the EIP-ring neurons, P-ring neurons or C-ring neurons. ***A***, The EIP-ring neurons expressed driver *c105-GAL4*, and the P-ring neurons expressed driver *VT5404-GAL4*. ***B***, The EIP-ring neurons expressed drivers *R31A12-GAL4* (Jenett et al., 2012) (the image is from Janelia Research Campus, https://flweb.janelia.org/cgi-bin/view_flew_imagery.cgi?line=R31A12#), *VT039763-GAL4* (Tirian and Dickson, 2017) (Janelia Research Campus, https://flweb.janelia.org/cgi-bin/view_flew_imagery.cgi?line=VT039763#), and *VT39763-GAL4*. ***C***, The P-ring neuron expressed control drivers *VT005404-GAL4* (Tirian and Dickson, 2017) (Janelia Research Campus, https://flweb.janelia.org/cgi-bin/view_flew_imagery.cgi?line=VT005404#) and *R14G09-GAL4* (Jenett et al., 2012) (Janelia Research Campus, https://flweb.janelia.org/cgi-bin/view_flew_imagery.cgi?line=R14G09#). ***D***, The C-ring neuron expressed driver *VT011965-GAL4* (Tirian and Dickson, 2017) (Janelia Research Campus, https://flweb.janelia.org/cgi-bin/view_flew_imagery.cgi?line=VT011965#). The Rubin lines *R31A12-GAL4*, *R14G09-GAL4* (Jenett et al., 2012) and the VT lines *VT039763-GAL4*, *VT005404-GAL4*, *VT011965-GAL4* (Tirian and Dickson, 2017) are from FlyLight database (https://www.janelia.org/project-team/flylight) and are provided under CC BY 4.0 license (https://reurl.cc/a5AD29). The drivers shown in ***B*-*D*** are less specific to the target neurons and are also expressed in many other brain regions. Therefore, they are not used in the present study.

**Extended Data Figure 1-2.**
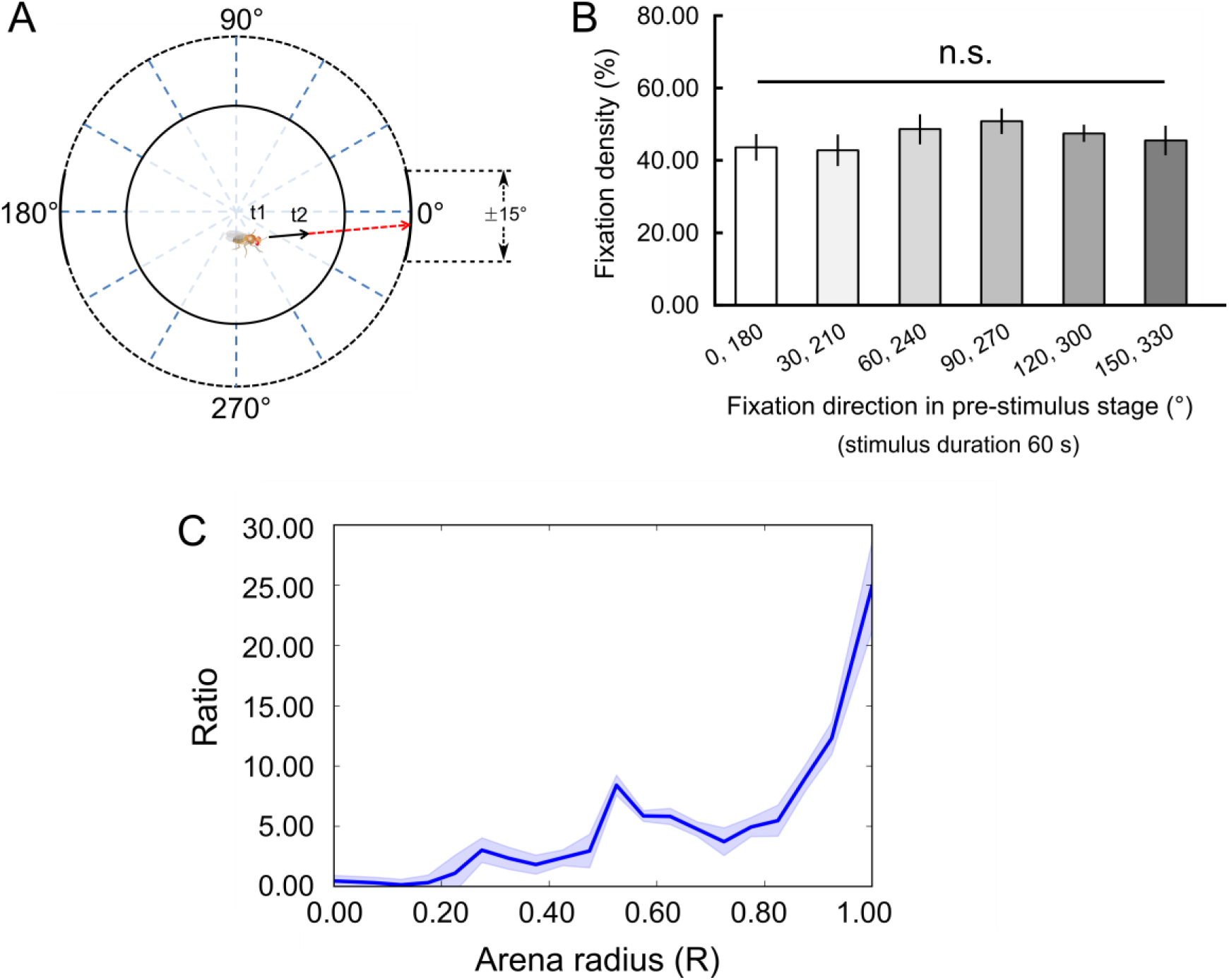
The definition of movement direction and the trajectory distribution. ***A***, In the behavioral task, we recorded the movement direction of the fly in each time step by calculating the vector (black arrow) that connects the positions of the fly in the previous (t1) and present (t2) time steps. The movement direction is defined by the projection (red arrow) of the vector on the LED screen. The movement direction is 356*°* (or -4*°*) in this case. ***B***, The population-averaged fixation density (fixation duration / stage duration) toward each pair of quantiles in the pre-stimulus stage. The fixation density of each pair of quantiles are comparable, suggesting that the flies did not exhibit directional preference in the first stage. ***C***, The distribution of the trajectories of flies on the platform along the radius during the pre-stimulus stage. The shaded area indicates the standard error of the mean. The values were normalized by the area spanned in each unit length of the radius.

**Extended Data Figure 1-3.**
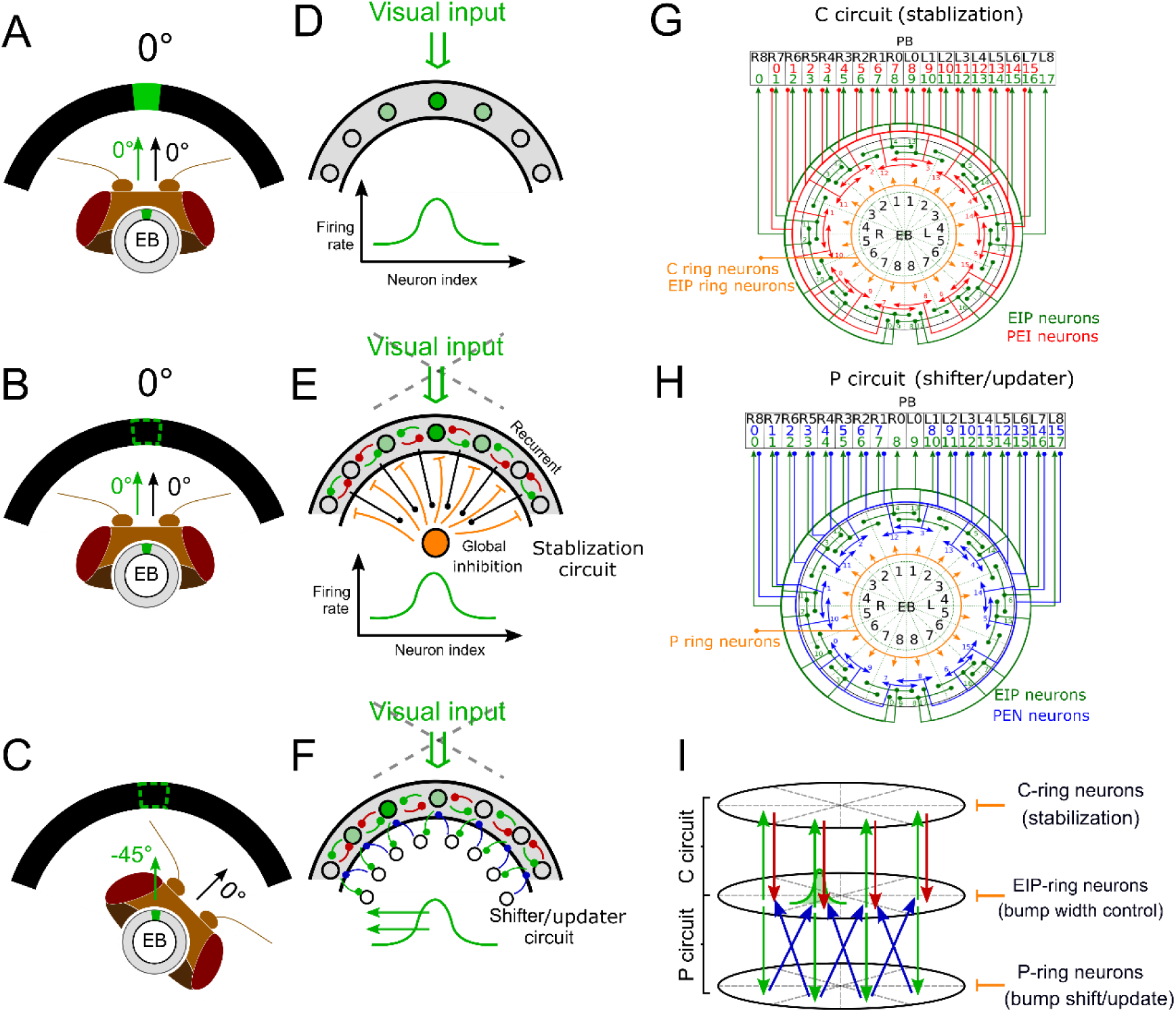
Schematics of the fruit fly compass circuits and its model. ***A,*** Fruit fly ellipsoid body (EB) (the grey torus) exhibits localized neural activity, termed activity bump (green), which corresponds to the direction of the visual cue on the screen. **B,** The activity bump persists when the visual cue is turned off. **C,** When the fruit fly changes its heading, the location of the activity bump shifts (or updates) so that it always indicates the head direction with respect to the cued direction. ***D***, The activity bump can be simply generated by localized input to neurons in EB. **E,** However, according to the attractor network model, the persistency of the bump after offset of the visual cue requires two types of synaptic connections. First, locally recurrent excitation (green and red), which creates the reverberatory activity in the absence of input. Second, the global feedback inhibition (orange), which controls the width of the activity bump so that it does not spread throughout the whole population of neurons. We call this two sets of connections “stabilization circuit”. **F,** Moreover, the shift of bump location due to the change of heading can be achieved by a shifter (or updater) circuit which form counterclockwise (blue, as shown) or clockwise (not shown) feedforward excitation, which propagates the activity in either direction. **G-H,** The specific connections as described in panels E and F were discovered in neurons that interconnect EB and PB (protocerebral bridge) in recent connectomic studies. The recurrent excitatory circuits are formed by the EIP (green) and PEI (red) neurons (C circuit), while the shifter circuits are formed by the EIP and PEN (blue) neurons (P circuit). These neurons are modulated by several types of GABAergic ring neurons (orange). The Su et al. 2017 model suggested that the coordinated activation of the C and P circuits are crucial for the function of spatial orientation. **I,** To simplify the visualization of the circuits, we redraw them using a three-ring representation, which highlights the roles of the ring neurons. The C-ring and P-ring neurons switch on/off the stabilization and shifter circuits, respectively. The EIP-ring neurons are responsible for controlling the width of the activity bump (the global inhibition in the attractor network). This representation is used in the subsequent figures in this paper. A-C, G & H are adapted from Su et al. Nat Comm 8, 139 (2017).

**Extended Data Figure 1-4.**
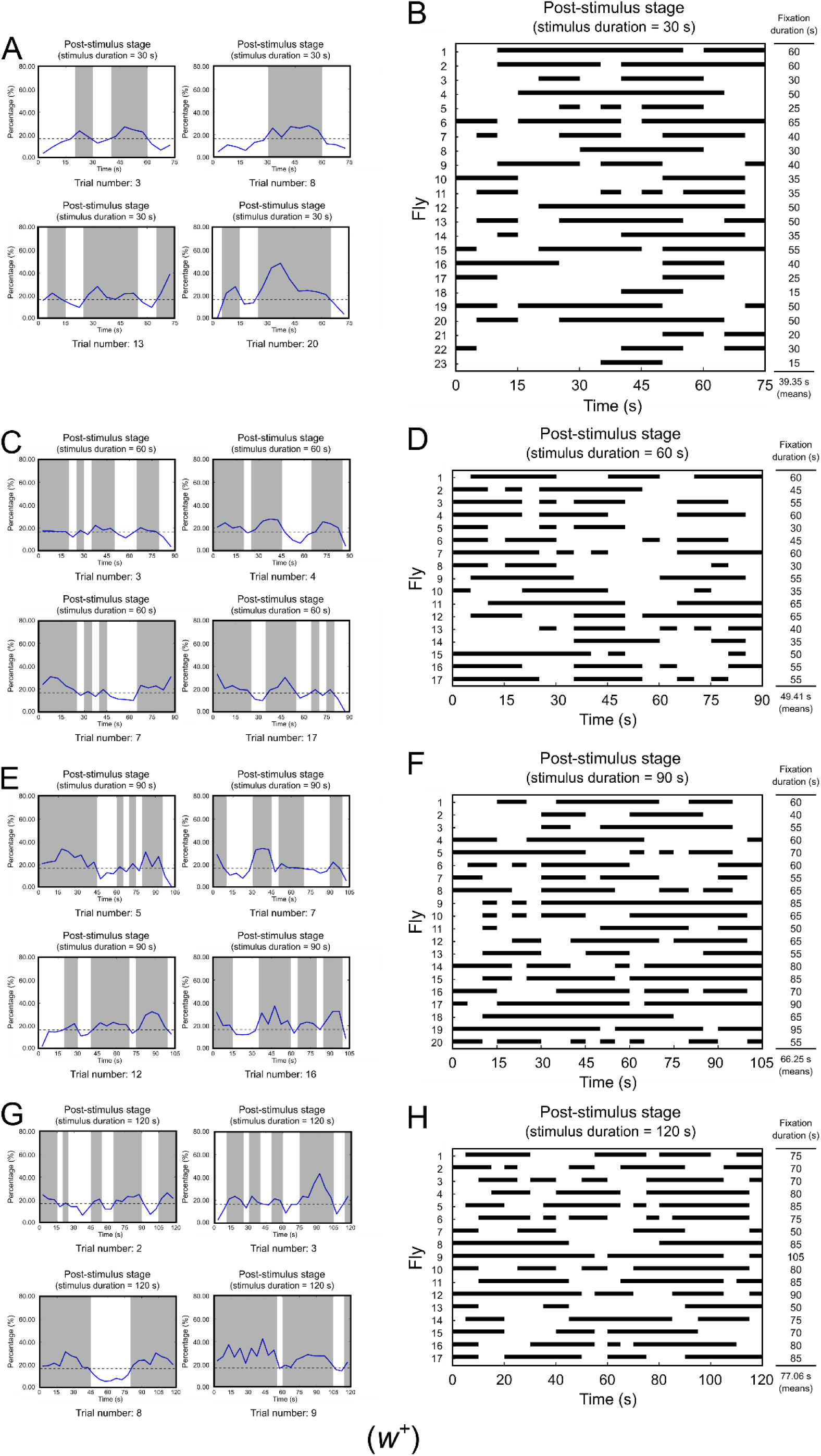
The movement patterns of Individual flies for difference stimulus stage durations. ***A***, Percentage of fixation for four example trials of wild-type flies. The shaded regions indicate the periods in which the percentage of movement toward the landmark positions is more than 16.67% (see Materials and Methods). ***B***, Distribution of the fixation bouts (black bars) in the post-stimulus stage for each wild-type flies (stimulus stage duration = 30 s). In this condition, the average fixation duration of each single trial is 39.35 s. ***C-D***, Same in ***A-B***, but with a stimulus duration of 60 s. In this condition, the average fixation duration of each single trial is 49.41 s. ***E-F***, Same in ***A-B***, but with a stimulus duration of 90 s. In this condition, the average fixation duration of each single trial is 66.25 s. ***G-H***, Same in ***A-B***, but with a stimulus duration of 120 s. In this condition, the average fixation duration of each single trial is 77.06 s.

**Extended Data Figure 2-1.**
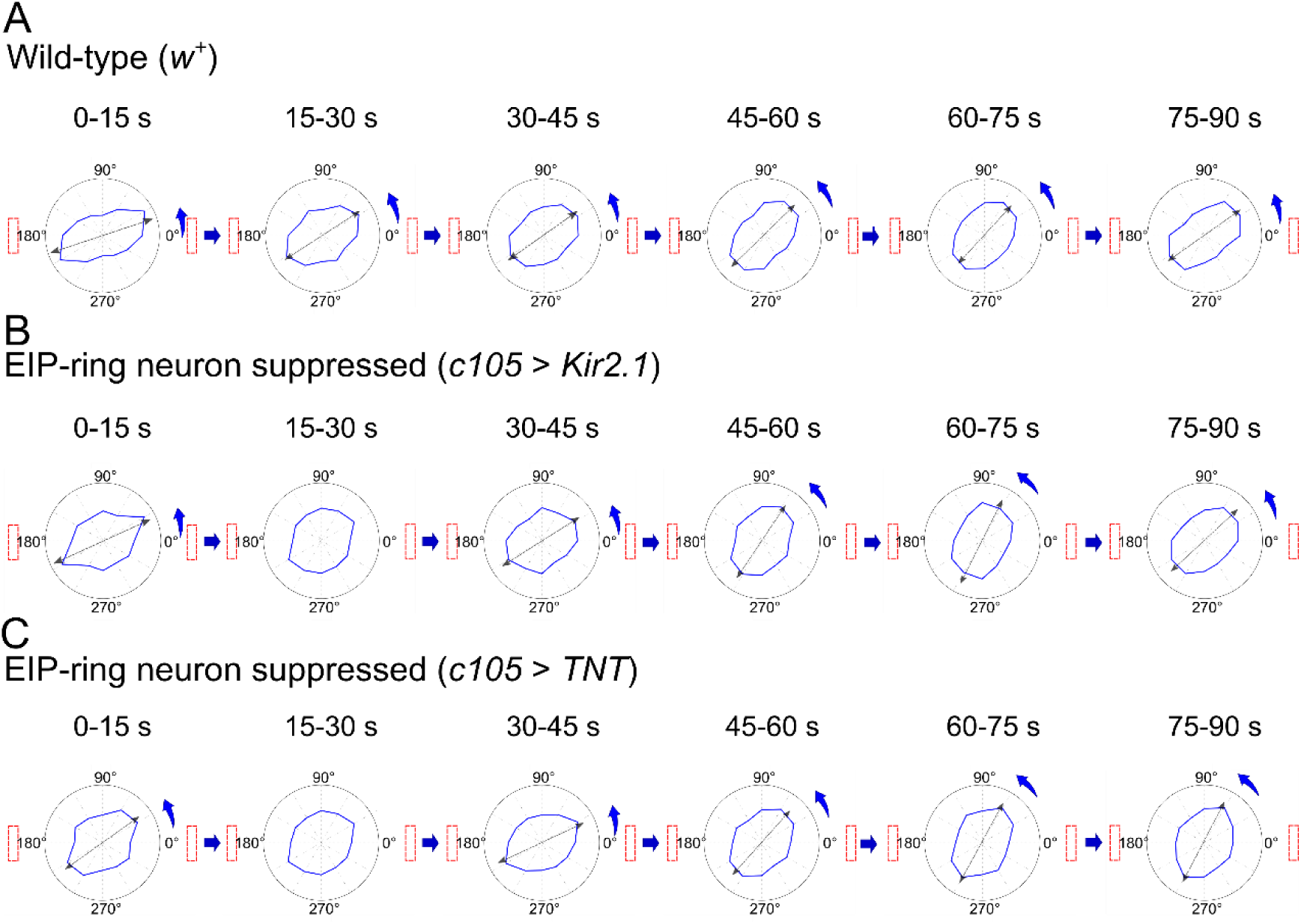
Radar plots of the flies in the control groups (wild-type: *w^+^*) and flies with suppressed EIP-ring neurons. ***A***, Wild-type *Drosophila* (genotype: *w^+^*). ***B-C***, Flies with suppressed EIP-ring neurons (***B*** for 32*°*C, *c105-GAL4, tub-GAL80^ts^;; UAS-Kir2.1* and ***C*** for 32*°*C, *c105-GAL4, tub-GAL80^ts^;; UAS-TNT*) in the post-stimulus stage.

**Extended Data Figure 3-1.**
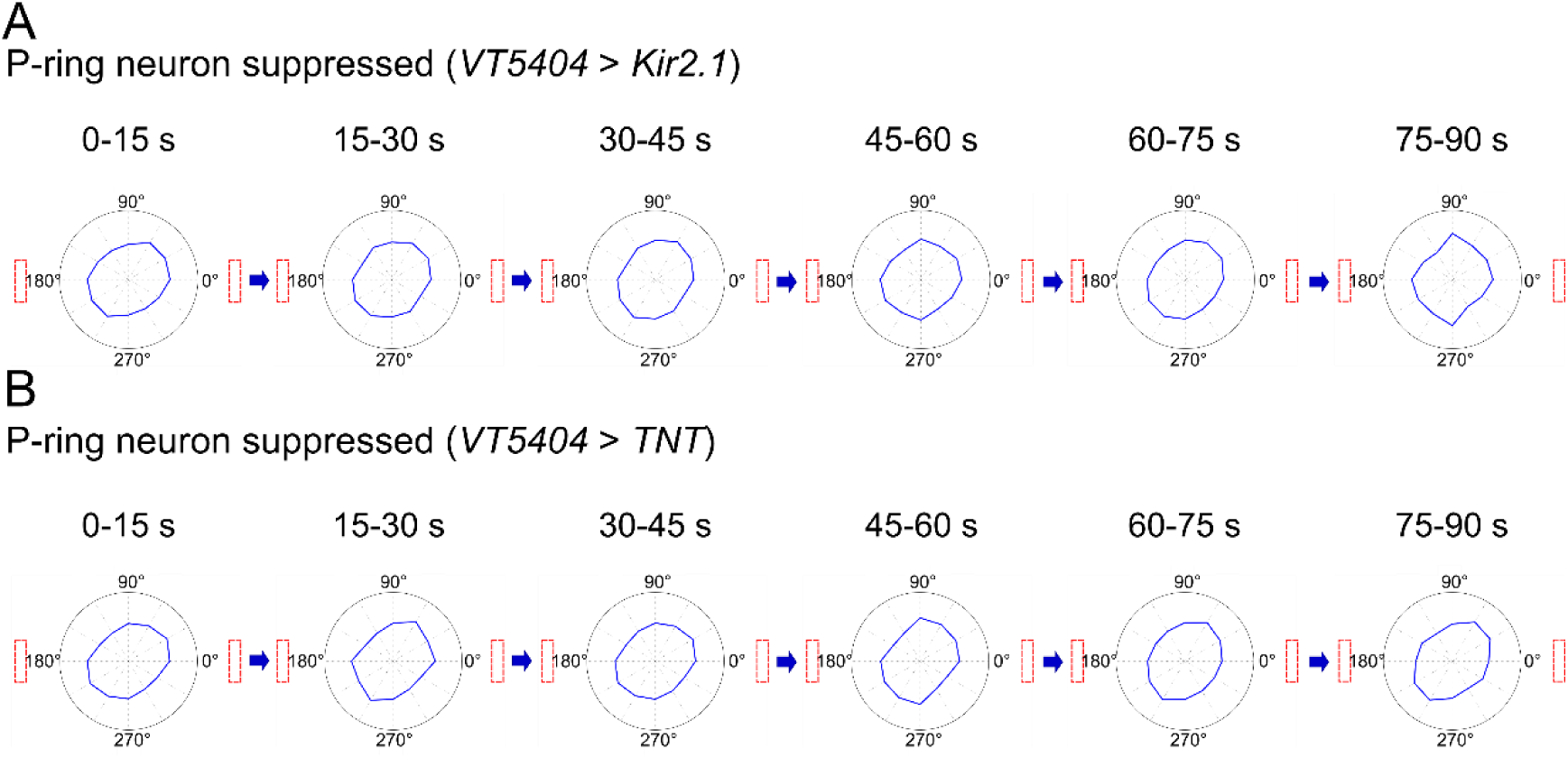
Radar plots of flies with suppressed P-ring neurons in the post-stimulus stage. ***A***, 32*°*C, *;;VT5404-GAL4, tub-GAL80^ts^/ UAS-Kir2.1*. ***B***, 32*°*C, *;;VT5404-GAL4, tub-GAL80^ts^/ UAS-TNT*.

**Extended Data Figure 4-1.**
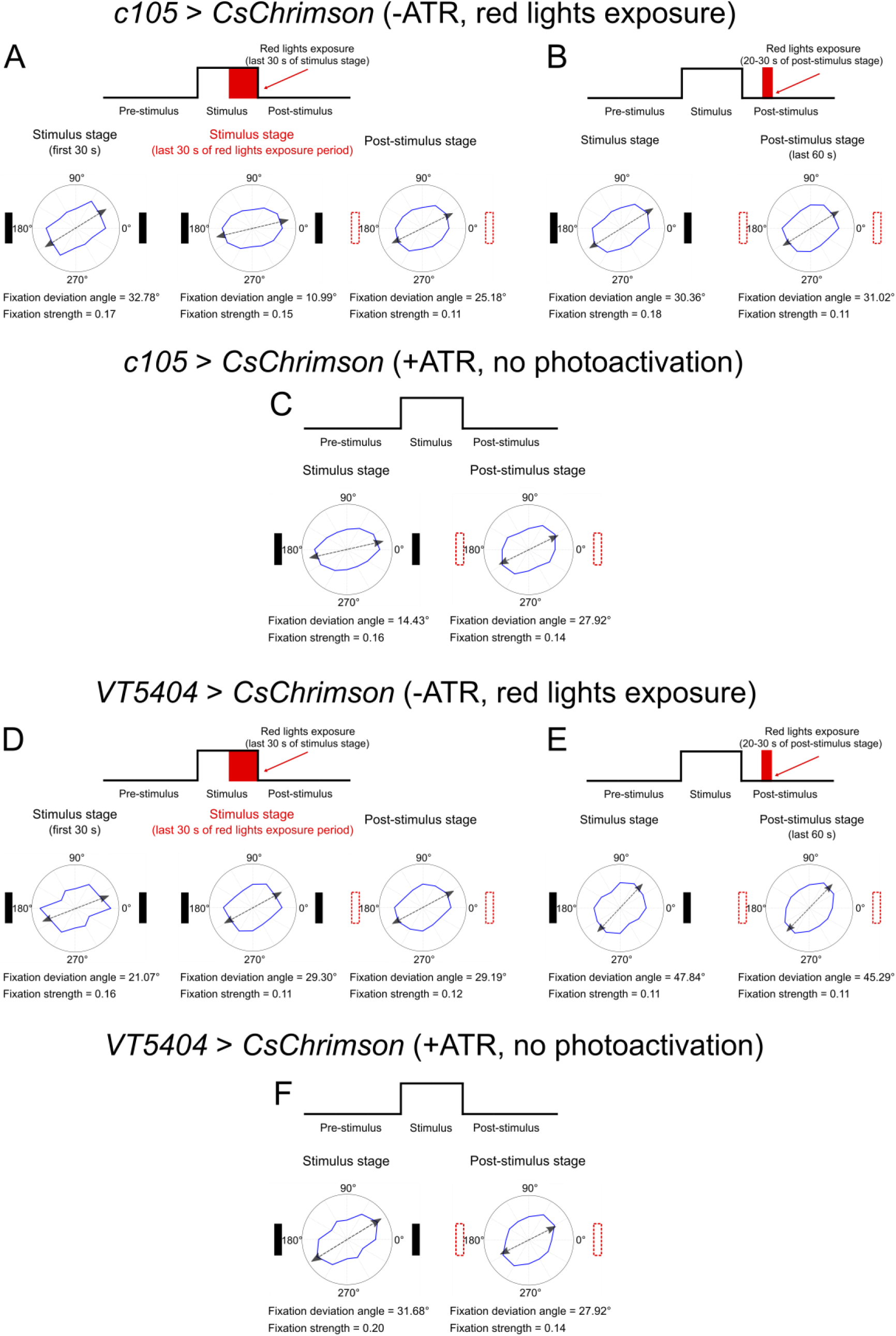
Performance of control groups with the same types of transgenic flies used in the optogenetic experiments. ***A-B***, Performance for flies that carry *c105-GAL4;;UAS-CsChrimson.mVenus*, but without feeding ATR. The schematic protocol (top) and the radar plots for the stimulus stage (bottom left, first 30s; bottom middle, last 30s) and the post-stimulus stage (bottom right). ***B***, Same as in ***A***, but with photoactivation during 20-30 s of the post-stimulus stage. ***C***, Same as in ***A***, but with ATR fed and no photoactivation. ***D-F***, Same as in ***A-C***, but for flies that carry *;;VT5404-GAL4/ UAS-CsChrimson.mVenus*. Both control groups exhibited a similar fixation performance to that of the wild-type flies.

**Extended Data Figure 4-2.**
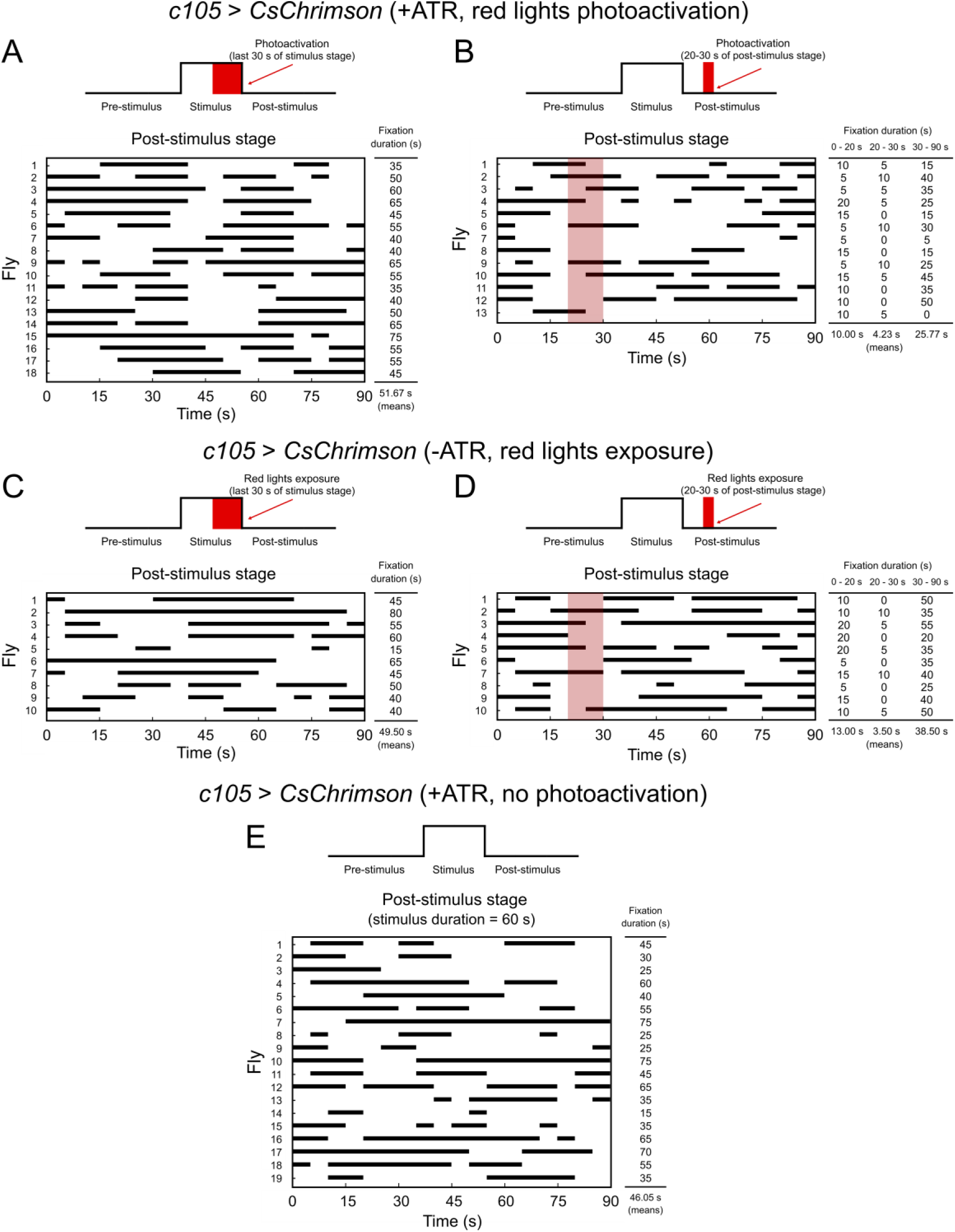
Movement patterns of individual flies in the photoactivation experiments. ***A***, Distribution of the fixation bouts (black bars) of each fly with the photoactivation of the EIP-ring neurons (*c105-GAL4;;UAS-CsChrimson.mVenus*) during the last 30 s of the stimulus stage. ***B***, Same as in ***A***. but with photoactivation during 20-30 s of the post-stimulus stage. ***C-D***, Same as in ***A-B***, but for non-ATR fed. ***E***, Same as in ***C***, but for ATR fed and no photoactivation was delivered.

**Extended Data Figure 4-3.**
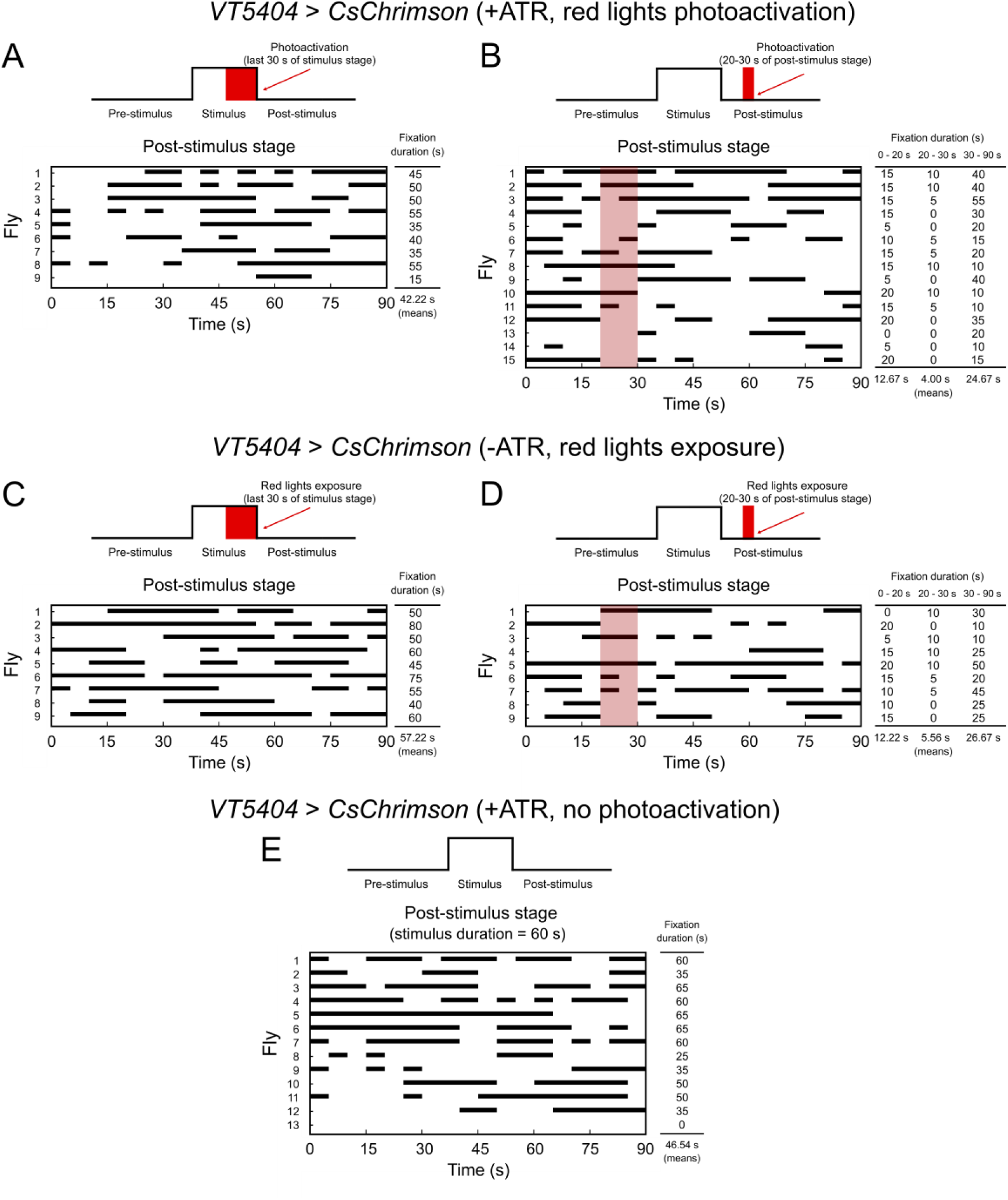
Same as in Extended Data Figure 4-2, but for flies that carry*;;VT5404-GAL4/ UAS-CsChrimson.mVenus*.

**Extended Data Figure 5-1.**
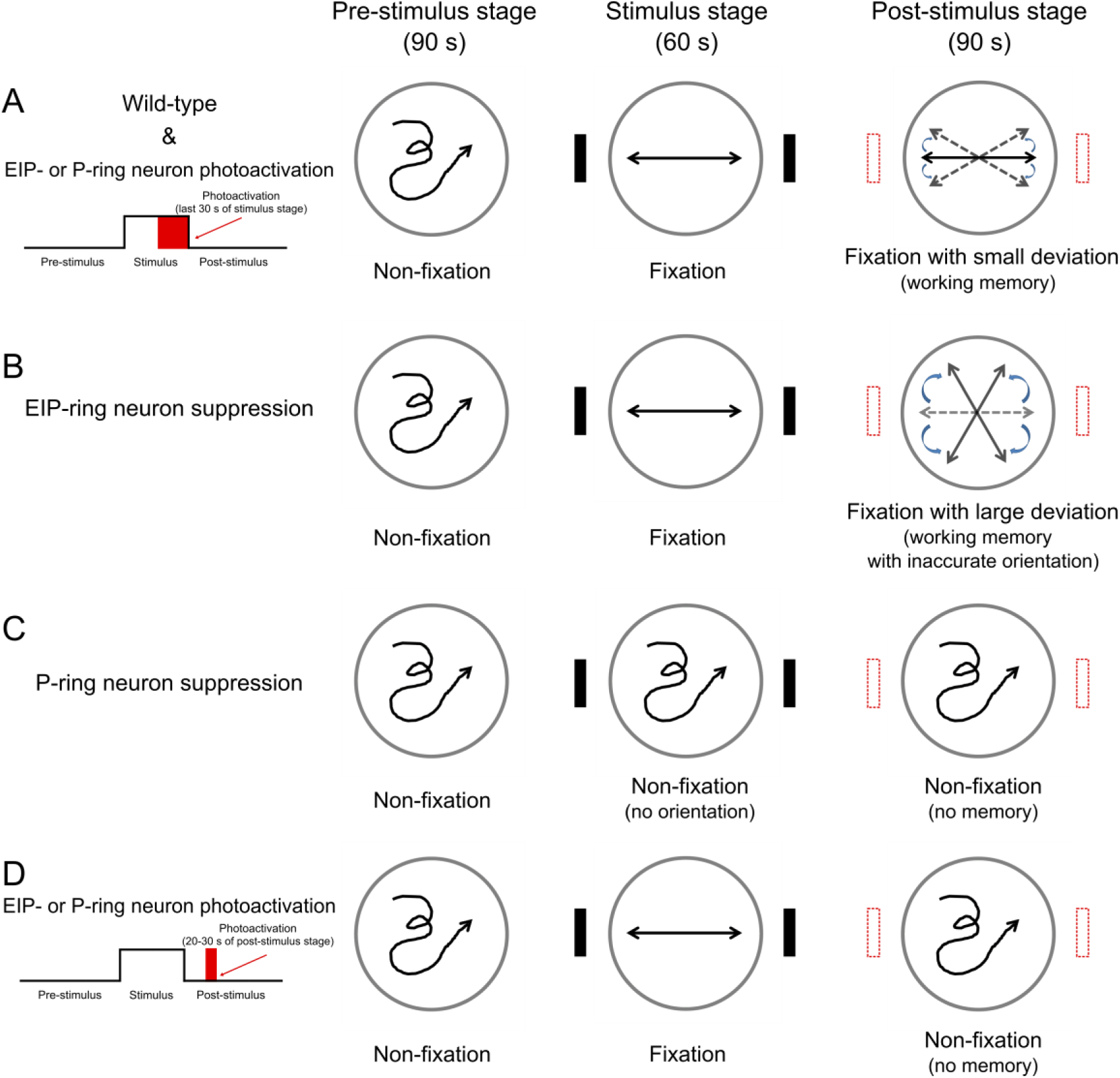
Schematic that summarizes the behavioral effects of EIP-ring and P-ring neuron suppression and photoactivation. ***A***, Wild-type flies maintain fixation behavior in both the stimulus stage and the post-stimulus stage, in which the flies exhibited a slight deviation in the fixation direction. Flies with transient photoactivation of the EIP-ring or P-ring neurons during the stimulus stage exhibited a similar behavioral pattern. ***B***, Flies with suppressed EIP-ring neurons exhibited fixation behavior with large deviation in the post-stimulus stages, respectively. ***C***, Flies with suppressed P-ring neurons could not maintain fixation behavior, indicating loss of spatial orientation. ***D***, Flies with transient photoactivation of the EIP-ring or P-ring neurons during the early post-stimulus stage eliminated the fixation behavior.

**Extended Data Figure 6-1.**
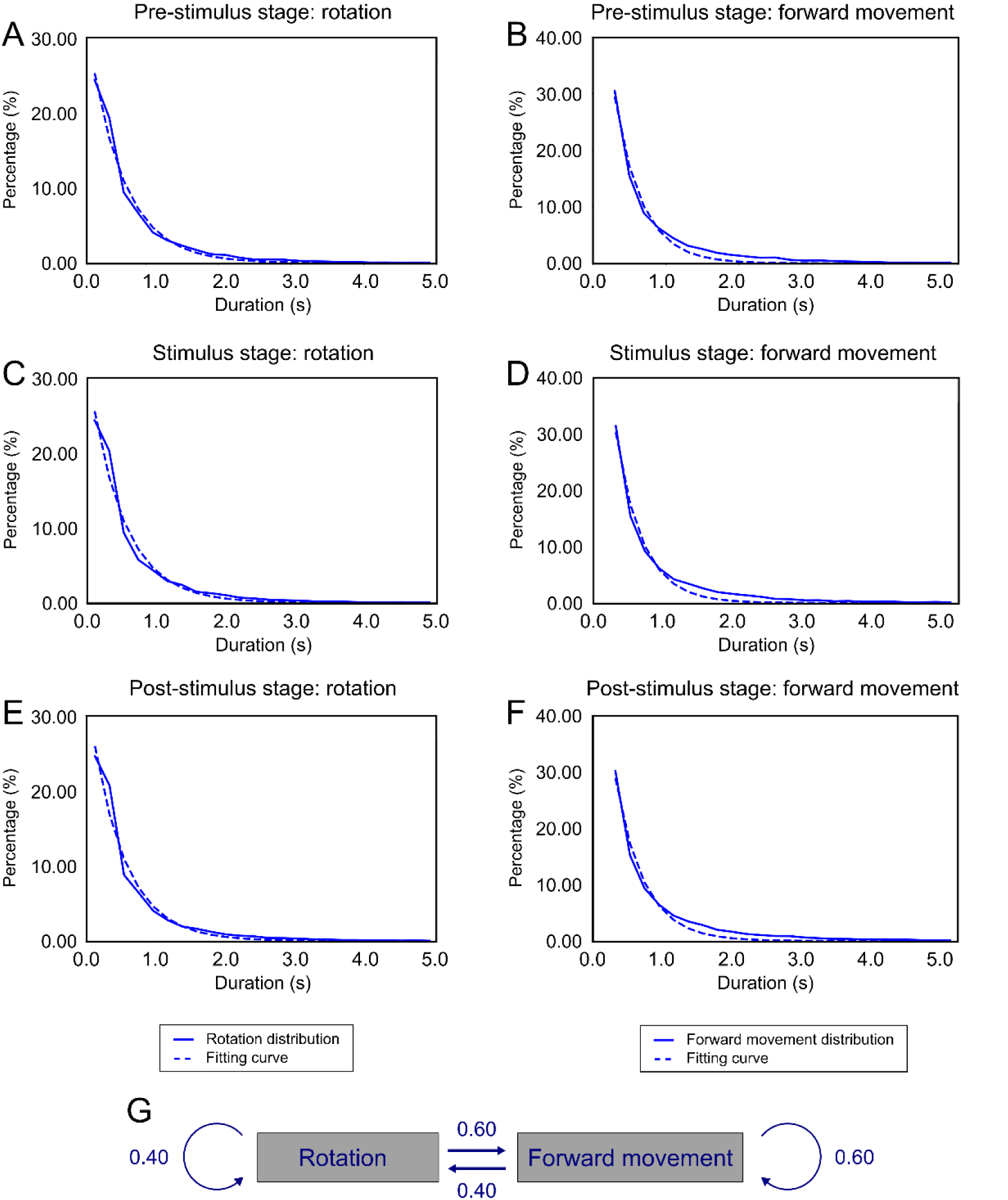
Distributions of duration of different behavioral states and the Markov chain model of behavioral control. Dashed lines represent the fitting curves, which is given by 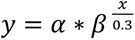. *α* is the fitting parameter and represents the probability of staying in the same state in each time window of 0.3 s. The exponential shape of the distribution of state duration indicates the Markov chain dynamics. ***A***, The distribution of duration of the rotation state in pre-stimulus stage. *α* = 0.473, *β* = 0.533 and *R*^2^ = 0.986. ***B***, Same as in ***A*** but for the forward movement state, *α* = 0.657, *β* = 0.448 and *R*^2^ = 0.985. ***C***, The distribution of duration of the rotation state in stimulus stage. *α* = 0.482, *β* = 0.529 and *R^2^* = 0.978. ***D***, Same as in ***C*** but for the forward movement state, *α* = 0.688, *β* = 0.439 and *R^2^* = 0.984. ***E***, The distribution of duration of the rotation state in post-stimulus stage. *α* = 0.495, *β* = 0.523 and *R^2^*= 0.975. ***F***, Same as in ***E*** but for the forward movement state, *α* = 0.621, *β* = 0.464 and *R*^2^ = 0.984. ***G***, The behavioral model was constructed based on the Markov chain dynamics and described the probability (given by the numbers next to the arrows) of a fly switching between behavioral states.

## Movie Legends

**Movie 1.** Laser escaping task for testing the visual perception of the flies with deficient photoreceptors.

**Movie 2.** Laser escaping task for testing the visual perception after amputation of foreleg in the wild-type flies.

**Movie 3.** Laser escaping task for testing the visual perception of the wild-type flies.

**Movie 4.** Laser escaping task for testing the visual perception of the flies with suppressed P-ring neurons.

**Movie 5.** A representative trial of EIP-ring neuron photoactivation during the last 30 s of the stimulus stage.

The landmarks located outside the movie and were placed at the left and right sides of the arena.

**Movie 6.** A representative trial of P-ring neuron photoactivation during the last 30 s of the stimulus stage.

The landmarks located outside the movie and were placed at the left and right sides of the arena.

